# Directional co-transcriptional folding and pausing create kinetic checkpoints for riboswitch-controlled gene expression

**DOI:** 10.64898/2026.03.06.710129

**Authors:** Adrien Chauvier, Javier Cabello-Villegas, Edward P. Nikonowicz, Nils G. Walter

**Affiliations:** Single Molecule Analysis Group and Center for RNA Biomedicine, Department of Chemistry, University of Michigan, Ann Arbor, Michigan, USA; Department of Biochemistry and Cell Biology, Rice University, Houston, TX 77005, USA

**Keywords:** Co-transcriptional RNA folding, ligand-sensing riboswitch, RNA polymerase, single molecule fluorescence microscopy, transcription

## Abstract

Transcriptional riboswitches typically regulate gene expression by sensing small metabolites or ions that cannot be fluorophore-labeled without perturbing recognition, limiting direct observation of ligand binding during transcription. The *GlyQS* T-box riboswitch instead senses the uncharged 3′ end of a macromolecular ligand, tRNA^Gly^, enabling visualization of co-transcriptional ligand binding. Using single-molecule fluorescence microscopy, we show that tRNA^Gly^ dynamically samples discrete structural domains of the nascent riboswitch as transcription proceeds. Productive recognition depends on a hierarchical 5′-to-3′ folding pathway shaped by transcription rate and pausing. Pausing enhances tRNA anchoring, stabilizes the riboswitch–tRNA complex, and selectively promotes RNA polymerase readthrough when an uncharged 3′ end is presented. These results support a kinetic checkpoint model in which early recruitment increases local ligand concentration, while pause-mediated stabilization converts transient encounters into committed regulatory decisions. More broadly, our findings identify directional co-transcriptional folding and transcriptional pausing as principles synergistically enabling selective and robust riboswitch function.

## Introduction

A defining feature of gene regulation mediated by non-coding RNAs is that it is intimately coupled to the progress of transcription, with the nascent RNA beginning to fold and adopt functional conformations as soon as it emerges from the RNA polymerase (RNAP)^1^. Co-transcriptional 5’-to-3′ folding imposes a dominant directional hierarchy to which RNAs have evolutionarily adapted, ensuring proper function in gene regulation^2,3^. In parallel, intrinsic pausing of RNAP at DNA-encoded sites and interactions with transcription factors have emerged as critical modulators of functional RNA folding, but the connections between co-transcriptional RNA folding and RNAP pausing remain underexplored^2,4^.

In bacteria, riboswitches are ubiquitous genetic elements, commonly found in the 5’-untranslated regions (UTRs) of messenger RNAs (mRNAs), where they respond to cellular metabolites or elemental ions to regulate gene expression at the level of transcription or translation^5^. Because they change conformation “on-the-fly,” riboswitches serve as tractable models for studying the role of transcription in shaping nascent RNA folding. Ligand sensing by the riboswitch aptamer domain has often been viewed as straightforward due to the presence of a well-conserved, robust ligand binding pocket^6^. However, recent studies on both transcriptional and translational riboswitches have revealed the coexistence of metastable structural intermediates that exchange on the timescale of transcription, defining a kinetic window for ligand binding and downstream expression platform manipulation^7–12^. This concept of a kinetically controlled structural partitioning is particularly evident in transcriptional riboswitches, where ligand recognition and any regulatory effects must occur before the RNAP reaches the termination point just downstream of the regulating RNA motif^12–15^.

T-box riboswitches are critical riboregulators, predominantly found in Gram-positive bacteria, where they control the expression of genes involved in amino acid synthesis, transport, and tRNA aminoacylation. This includes the archetypical *Bacillus subtilis (Bsu-)GlyQS* T-box riboswitch, which regulates the *GlyQS* operon encoding the glycyl-tRNA synthetase enzyme^16^. Each T-box binds a specific tRNA, sensing the uncharged 3′-aminoacylation state as a metabolic signal that guides gene expression^17^. A typical transcriptional T-box riboswitch is modular and consists of two main domains: a 5’-decoding or aptamer domain (including Stems I, II, and IIA/B) and a 3′-discriminator domain (including Stem III and the competing transcriptional Antiterminator, AT, and Terminator, T, hairpins), connected by a single-stranded linker^17^. The decoding domain binds the tRNA through its Stem I, while the discriminator domain senses the charging state of the tRNA 3′-NCCA end, shifting its conformation typically from the T to the AT hairpin to allow downstream transcription. While the decoding region exhibits variability, the overall domain architecture is highly conserved across bacterial species^18^.

Recent structural advances through X-ray crystallography^19–21^ and cryo-electron microscopy^22^ have provided detailed, yet static views of the T-box riboswitch core in complex with tRNA ligands, highlighting three interaction sites for the anticodon, elbow and 3′-NCCA end (Fig. 1a). These studies have clarified how specific contacts stabilize high-affinity tRNA binding and uncharged 3′-end recognition. Nevertheless, the dynamic conformational changes underlying tRNA sensing―particularly during the continuous release of 3′-sequence by the transcribing RNAP―have remain enigmatic. Recent single-molecule fluorescence resonance energy transfer (smFRET) studies of defined RNA-only complexes^23–26^ suggest that tRNA engagement occurs via a hierarchical two-step mechanism that adjusts the riboswitch conformation, with implications for gene regulation^27^. Despite these advances, the real-time dynamics of ligand binding during transcription have remained elusive.

**Fig 1.**
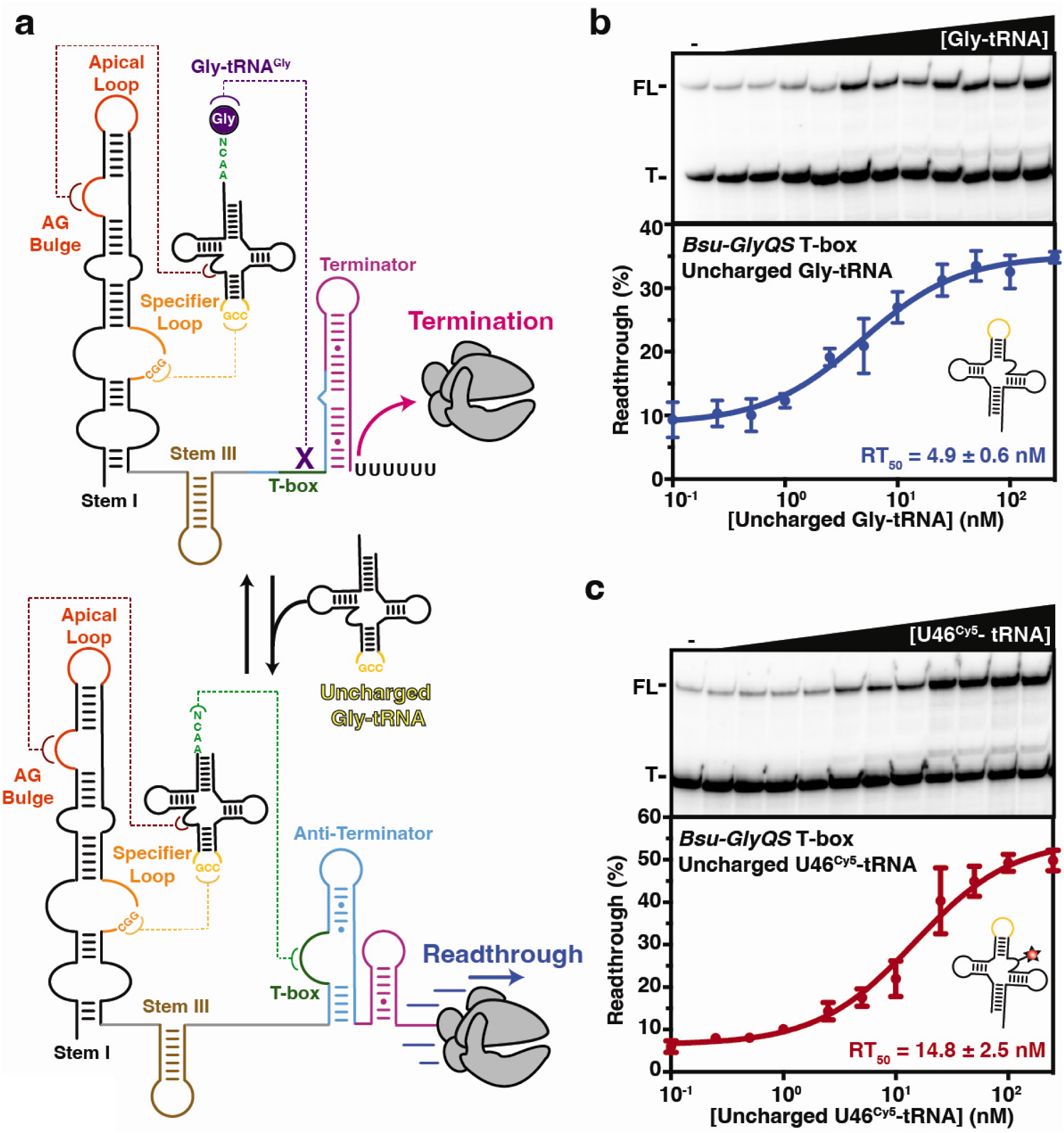
Transcriptional Regulation Induced by the *Bsu-GlyQS* T-box Riboswitch. **a**, The *GlyQS* T-box riboswitch from *B. subtilis* regulates gene expression at the transcriptional level. Stabilization of the Antiterminator (AT) upon binding of the uncharged tRNA^Gly^ prevents the formation of an intrinsic Terminator (T) hairpin leading to the transcription of the downstream gene. tRNA selection is performed through interaction between the Specifier Loop within Stem I and the anticodon of the tRNA. Steric hindrance caused by the presence of glycine amino acid (charged tRNA – Gly-tRNA^Gly^) prevents the interaction between the tRNA 3′-end and the T-box motif which would otherwise promotes the formation of the AT. **b-c**, Gly-tRNA (**b**) and U46^Cy5^-tRNA (**c**) dependent single round transcription assay of a DNA template containing the 5’-UTR of the *GlyQS* operon from *B. subtilis.* Transcription reactions were performed using the *E. coli* RNAP with 10 µM rNTPs. ^32^P-labeled products were resolved on 6% polyacrylamide denaturing gel separating the full-length (FL) and terminated (T) RNA products. Plot of the fraction of transcriptional readthrough versus the concentration of tRNA ligand are displayed below the representative gels. RT_50_ are indicated. Error bars represent the Standard Deviation (SD) of the mean from independent replicates.

Beyond their mechanistic intrigue, and given their prevalence in pathogenic bacteria, T-box riboswitches have garnered attention as potential antimicrobial targets^18,28,29^. Elucidating the detailed mechanisms by which T-box riboswitches sense and respond to tRNA binding thus promises not only to advance our understanding of bacterial gene regulation but also to inform the development of novel antimicrobial strategies^30,31^. Together, these motivations call for the development of tools that reveal the temporal coordination of ligand recognition and structural rearrangements of the elongating T-box sequence.

In this study, we exploited the substantial size of the tRNA ligand to site-specifically introduce a Cy5 label at U46 with minimal functional disruption, generating U46^Cy5^-tRNA, and used single-molecule fluorescence microscopy to dissect the dynamic co-transcriptional mechanism by which the *Bsu*-GlyQS T-box riboswitch activates gene expression. Our findings highlight the critical role of 5’-to-3′ directional T-box RNA folding, which we find modulated by intrinsic RNAP pausing, for efficient tRNA^Gly^ sensing during transcription. Using the corresponding paused elongation complexes (PECs) together with *in situ* single-molecule probing approaches, we demonstrate that RNAP proximity enhances tRNA recruitment, facilitated by strategic positioning of the pauses. AT folding was found to expose the T-box motif, promoting a stabilized co-transcriptional capture of tRNA^Gly^. Kinetic analysis of ligand binding further supports a hierarchical mechanism in which early, transient tRNA^Gly^ interactions with the first-transcribed Stem I (“sensing”) are crucial for subsequent 3′-NCCA end capture (“anchoring”) by the emerging AT. Moreover, by modulating transcription rates, we establish that RNAP speed constrains ligand recognition within a variable, but narrow kinetic window. Finally, deploying a double-PIFE (protein-induced fluorescence enhancement) approach, we link stabilized tRNA^Gly^ anchoring to transcriptional readthrough under near-physiological conditions, unmasking a temporally orchestrated regulatory mechanism. Together, these findings provide a comprehensive understanding of how the *Bsu-GlyQS* T-box riboswitch integrates dynamic, directional RNA folding and RNAP pausing with a hierarchical ligand sensing mechanism to regulate essential metabolic pathways―as a paradigm for riboswitch mechanisms more broadly and a steppingstone for antimicrobial interference with T-box function.

## Results

### Probing co-transcriptional *Bsu-GlyQS* T-box function using fluorescently labeled tRNA

The *Bsu*-*GlyQS* T-box riboswitch senses the uncharged tRNA^Gly^ to promote the transcription of the downstream *GlyQS* operon encoding glycyl-tRNA synthetase^29,30^. As expected, single-round *in vitro* transcription of the *Bsu*-*GlyQS* T-box riboswitch displays efficient transcription readthrough in the presence of uncharged tRNA (Fig. 1b). In accordance with previous studies^30^, we obtained an RT_50_ (i.e., the concentration of tRNA^Gly^ needed to induce 50% of transcription readthrough regulation) of 4.9 ± 0.6 nM, showing efficient co-transcriptional ligand sensing and gene regulation upon tRNA^Gly^ binding (Fig. 1b).

To monitor how the *Bsu*-*GlyQS* T-box riboswitch senses its cognate ligand during transcription, we engineered a fluorescently labeled tRNA^Gly^ (Supplementary Fig. 1a,b)^25^, assembled by ligating two synthetic RNA strands to incorporate a Cy5 fluorophore at position U46, within the tRNA elbow (U46^Cy5^-tRNA)—far from the functionally critical 3′-NCCA end. Using this labeled uncharged tRNA during single-round *in vitro* transcription assays, we obtained an RT_50_ of 7.4 ± 2.5 nM (Fig. 1c). This only modestly elevated value may reflect a minor perturbation of the tertiary interaction between the dual T-loop motif (DTM) in Stem I and the elbow region of the labeled tRNA^Gly^. By comparison, a ΔDTM mutant, which entirely disrupts this tertiary contact^25^, showed a substantially (∼16-fold) higher RT_50_ of 80 ± 7.6 nM (Supplementary Fig. 1c), highlighting the full extent of the DTM’s contribution to tRNA binding while underscoring that perturbation by fluorophore labeling is minor.

Together, our results demonstrate that U46^Cy5^-tRNA has robust functionality in promoting transcriptional readthrough, validating its suitability for single-molecule fluorescence studies. Unlike typical riboswitches that encapsulate their small-molecule or elemental ion effector within a tight binding pocket so that they cannot be fluorophore labeled, the T-box riboswitch model system thus offers a unique opportunity to introduce a fluorophore label and directly interrogate the dynamics of co-transcriptional ligand interactions at the single-molecule level.

### Proximal RNAP stabilizes tRNA anchoring to two programmed paused elongation complexes

Given the indirect evidence available for other transcriptional riboswitches for which ligand binding could not be observed directly^13,14,32,33^, co-transcriptional tRNA sensing is expected to be required to efficiently promote transcription readthrough before the RNAP reaches the termination point. To measure the kinetics of ligand binding at functionally defined stages of transcription, we first performed transcription assays at limiting rNTPs concentration, confirming previously reported^34,35^ RNAP pause sites along the *Bsu*-*GlyQS* T-box transcript at positions 138, 178 and 207 (Supplementary Fig. 1d,e). We then assembled Paused Elongation Complexes (PECs) from the downstream biotinylated, streptavidin-capped DNA template, RNAP, and Cy3-labeled *Bsu*-*GlyQS* T-box RNA that was generated and folded by Cy3-primed transcription in situ^11^. We immobilized this complex on a biotin-PEG coated microscope slide^14^, and monitored U46^Cy5^-tRNA binding events by colocalizing the Cy5 and Cy3 fluorescence signals in a total internal reflection fluorescence (TIRF) microscope (Fig. 2a-c). Consistent with earlier findings that used polymerase-free, isolated T-box RNA fragments of varying length^25^, U46^Cy5^-tRNA binding in the presence of RNAP is transient and dynamic, emphasizing the need for real-time single-molecule measurements to capture these events during the co-transcriptional sensing process.

**Fig 2.**
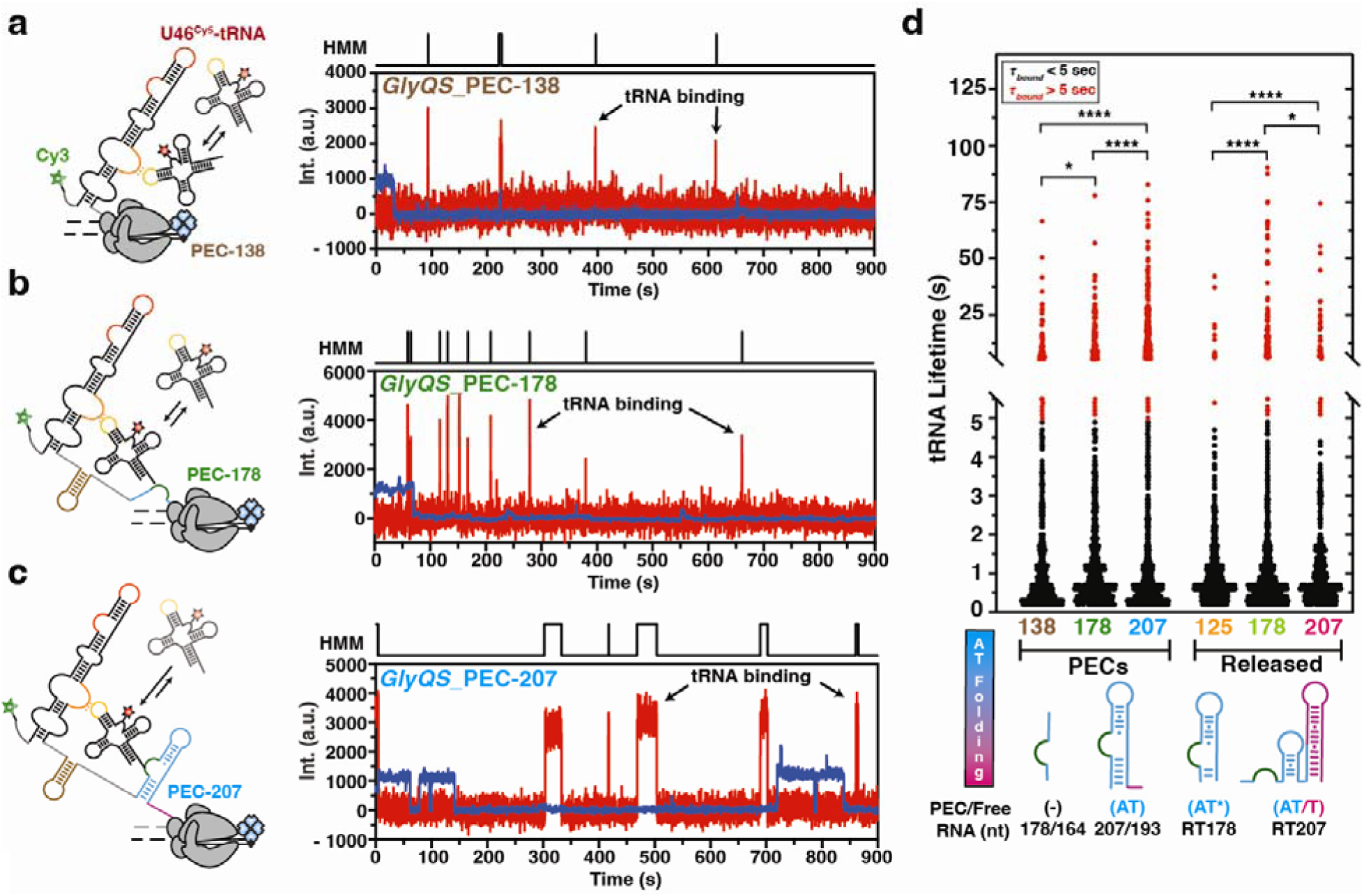
Monitoring tRNA^Gly^ Binding Kinetics at Key Transcriptional Pause Sites. **a-c**, Secondary structure of the *Bsu-GlyQS* T-box riboswitch in the context of the Paused Elongation Complexes (PEC) at the 138 (**a**), 178 (**b**) and 207 (**c**) positions, alongside representative single-molecule trajectories showing U46^Cy5^-tRNA binding (red) to the corresponding PEC (blue). Hidden Markov Modeling (HMM) is indicated on the top of each trace. **d**, Plot displaying the overall distribution of tRNA lifetimes for the different constructs tested. Short binding events (τ*_bound_* <5 s) are indicated in black and long-lived binding events (τ*_bound_* >5 s) are indicated in red. Predicted secondary structures of the AT is shown at the bottom with the corresponding length of the construct and the free RNA emerging from the RNA exit channel in the case of PECs. The total number of molecules analyzed is as follows: PEC-138 = 204; PEC-178 = 208; PEC-207 = 195; RT-125 = 236; RT-178 = 237; RT-207 = 203. (****P < 0.001, ***P < 0.005, **P < 0.01, *P < 0.05)

For the early transcription intermediate PEC-138, which presents only Stem I outside the RNAP’s exit channel, most U46^Cy5^-tRNA^Gly^-T-box interactions were short-lived: 90% dissociated within ∼1 s and only a few persisting beyond 5 s (Fig. 2a and d). As *Bsu*-*GlyQS* transcription advanced to PEC-178 (adding the 5’ segment of the T-box motif) and PEC-207 (adding the full T-box motif), the fraction of longer-lived binding events increased to ∼12% and 25%, respectively, resulting in overall (weighted-average) decreasing dissociation rate constants of 0.40 (PEC-138), 0.28 (PEC-178) and 0.17 s^-1^ (PEC-207; Fig. 2c,d and Supplementary Table 2). Concurrently, the overall binding rate constant (k_on_) increased from 0.75 × 10^6^ M^-1^ s^-1^ at PEC-138 to 0.94 × 10^6^ M^-1^ s^-1^ at PEC-178 to 1.25 × 10^6^ M^-1^ s^-1^ at PEC-207 (Supplementary Table 2). These results support a model in which emergence of the downstream expression platform both accelerates local ligand recruitment and stabilizes the riboswitch-tRNA complex once formed.

To test whether the tRNA 3′ end affects recognition, we compared PEC-207 binding of unmodified U46^Cy5^-tRNA^Gly^ with tRNAs bearing two 3′-blocking modifications—either glycyl aminoacylation or a 3′ Cy5 label. These modified tRNAs bound much less efficiently, with the overall association rate constant decreasing to 0.48 × 10^6^ M^-1^ s^-1^ (glycyl) and 0.32 × 10^6^ M^-1^ s^-1^ (Cy5; Supplementary Fig. 4a–d). However, once complexes were formed, their lifetimes and the fraction of long-lived events remained essentially unchanged (0.14 s^-1^ for glycyl and 0.21 s^-1^ for Cy5) relative to uncharged U46^Cy5^-tRNA^Gly^ (0.17 s^-1^; Supplementary Fig. 4e), indicating that aminoacylation impairs association but does not destabilize already formed complexes. Furthermore, pre-incubation of PEC-207 with a high uncharged tRNA concentration (250 nM) revealed the capacity to form ultra-stable complexes (mean bound time ∼355 s; Supplementary Fig. 3a–b), comparable to a previously report for an RNAP-free, isolated T-box construct of similar size^25^. Together, these data indicate that steric obstruction of the 3′ end (by tRNA charging or labeling) reduces sensing by lowering association efficiency rather than by shortening the complex’s lifetime.

To quantify how the presence of RNAP alters ligand-sensing kinetics, we compared U46^Cy5^-tRNA^Gly^ binding to PECs with binding to the corresponding released transcripts (RTs)—transcripts that were allowed to fold co-transcriptionally, but then released from the transcription bubble, and immobilized directly onto the TIRF surface (Fig. 2d). For RT-125, which contains only Stem I and lacks RNAP, tRNA binding closely resembles that observed for the corresponding PEC-138, indicating a minimal RNAP effect at this early pause. In contrast, the weighted-average dissociation rate constant is substantially higher for RT-125 (1.08 s^-1^) compared to PEC-138 (0.40 s^-1^), suggesting a stabilizing effect of RNAP during early transcription (Fig. 2c, d and Supplementary Table 2). Similarly, RT-207, which encompasses the full T-box motif but again is free of RNAP, showed a marked drop in the fraction of long-lived U46^Cy5^-tRNA events (∼7%) compared with PEC-207 (∼25%), leading to a substantial ∼3-fold decrease of the weighted-average dissociation rate constant in the presence of RNAP (0.53 s^-1^ for RT-207 versus 0.17 s^-1^ for PEC-207) (Fig. 2c, d and Supplementary Table 2). Finally, RT-178 exhibited the highest tRNA complex stability among the released transcripts (Fig. 2d), presumably due to the absence of the complementary 3′ segment of the AT hairpin.

These differences are consistent with a model in which RNAP sterically restricts folding of the 3′-most ∼10-14 nucleotides that reside in the RNA–DNA hybrid and RNAP exit channel^36^, thereby delaying formation of the AT and T hairpins in PEC-178 and PEC-207 and, as a consequence, promoting the T-box:tRNA 3′-end interaction (Fig. 2d). Supporting this idea, RT-207 forms ultra-stable complexes with a mean bound time (∼256 s) that is ∼28% shorter than that measured for PEC-207 (Supplementary Fig. 3c,d), further indicating that completion of T-hairpin folding (permitted only when RNAP is absent) competes with and weakens tRNA 3′-end engagement by the Bsu-GlyQS T-box.

These data indicate that transcriptional pausing actively guides co-transcriptional folding and ligand recognition by delaying or preventing premature formation of competing 3′-end structures. Co-transcriptionally folded RTs recruit U46^Cy5^-tRNA efficiently (Supplementary Fig. 2e,f), and their binding rate constants mirror the trend observed for paused elongation complexes. Conversely, the proximal RNAP—specifically at pauses programmed at positions 178 and 207—enhances tRNA-anchoring stability during transcription by suppressing competing intrinsic T-box structures.

### Folding of the antiterminator promotes stable tRNA anchoring

To identify the key structural determinants required for stable co-transcriptional tRNA anchoring, we introduced mutations in the T-box motif (mutant Mbox) and the AT hairpin (MAT) (Supplementary Fig. 1a). These mutations were incorporated into PEC-207, which binds U46^Cy5^-tRNA most efficiently and features the fully transcribed AT domain, while minimizing potential interference from the thermodynamically more stable T hairpin (Fig. 3).

**Fig 3.**
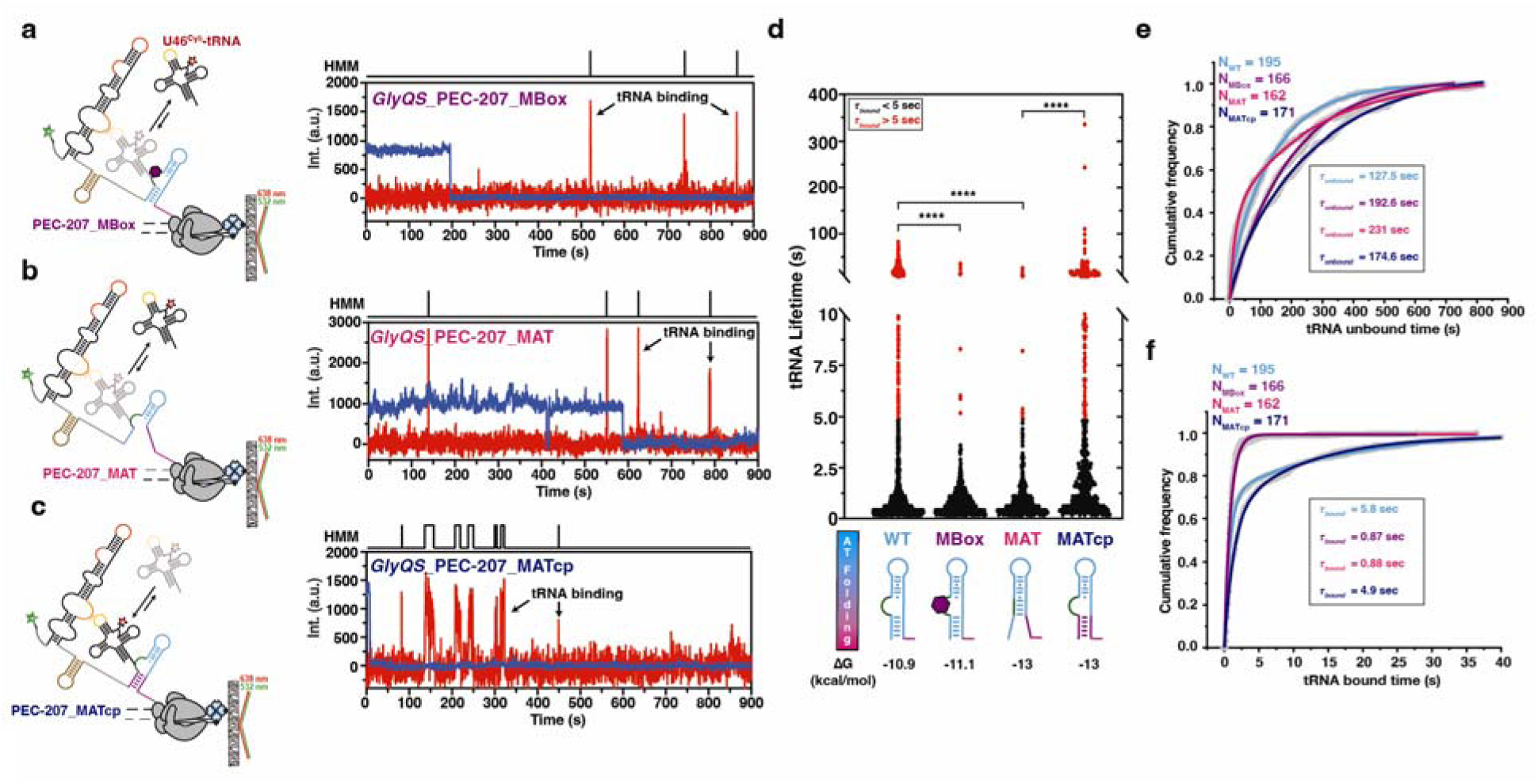
Folding of the Antiterminator Promotes Ligand Sensing and 3′-end anchoring. **a-c**, Secondary structure of the *Bsu-GlyQS* T-box riboswitch in the context of the PEC-207_MBox (**a**), PEC-207_MAT (**b**) and PEC-207_MATcp (**c**), alongside representative single-molecule trajectories showing U46^Cy5^-tRNA binding (red) to the corresponding PEC (blue). Hidden Markov Modeling (HMM) is indicated on the top of each trace. **d**, Plot displaying the overall distribution of tRNA lifetimes for the different constructs tested. Short binding events (τ*_bound_* <5 s) are indicated in black and long-lived binding events (τ*_bound_* >5 s) are indicated in red.. Predicted secondary structure of the AT is shown at the bottom with the calculated ΔG from UNAfold^56^. The total number of molecules analyzed is as follow: PEC-207_WT = 195; PEC-207_Mbox = 166; PEC-207_MAT = 162; PEC-207_MATcp = 171; (****P < 0.001, ***P < 0.005, **P < 0.01, *P < 0.05). **e-f**, Plots displaying the cumulative unbound (**e**) and bound (**f**) dwell times of U46^Cy5^-tRNA in the different PEC-207 constructs. The overall unbound (τ*_unbound_)* and bound (τ*_bound_)* times are indicated.

As expected, disrupting the tRNA 3′-end recognition site (Mbox) substantially impaired U46^Cy5^-tRNA anchoring, yielding only short-lived binding events and an overall dissociation rate constant comparable to that observed for Stem I only (1.16 s^-1^ for PEC-207-Mbox) (Fig. 3a, d and Supplementary Table 2). Accordingly, *in vitro* transcription assays using the Mbox mutation showed no substantial effect of tRNA^Gly^ on transcription termination (Supplementary Fig. 1f), reinforcing the essential role of this interaction in ligand-mediated gene regulation.

Strikingly, the PEC-207-MAT mutant exhibited binding behavior similar to PEC-207-MBox, displaying only transient U46^Cy5^-tRNA binding with short bound times (Fig. 3b, d and f). This finding suggests that proper AT folding is essential for suitably presenting the T-box motif, thereby enabling stable co-transcriptional tRNA anchoring. Similarly, both mutants showed U46^Cy5^-tRNA unbound times comparable to the Stem I-only RNA (PEC-138; Fig. 3e and Supplementary Table 2), underscoring the importance of both T-box motif and AT hairpin also in initial tRNA^Gly^ recognition. Notably, the introduced mutations reside within the switching region that participates in both AT and T structures in the full-length transcript^20,34^ (Supplementary Fig. 1a), indicating that alternating structures in this region could affect tRNA^Gly^ anchoring during transcription.

To further test this folding mechanism, we introduced compensatory mutations to restore the base pairing at the bottom of the AT stem, stabilizing its predicted secondary structure (MATcp; Supplementary Fig. 1a). In the context of the resulting PEC-207-MATcp, both short-and long-lived U46^Cy5^-tRNA binding events were recovered, further supporting the idea that AT folding promotes T-box motif display for stable co-transcriptional tRNA anchoring (Fig. 3c, d and f)..

Together, our kinetic and mutational analyses reveal that AT folding promotes efficient co-transcriptional tRNA^Gly^ sensing by presenting the T-box motif for stable tRNA 3′-end recognition.

### Real-time monitoring reveals sequential co-transcriptional recruitment and anchoring of tRNA

Next, to understand when tRNA sensing and capture occur during active transcription, we employed a single-molecule assay that monitors both transcription and U46^Cy5^-tRNA sensing in real-time^14,37,38^. To this end, DNA templates encoding either Stem I only (*GlyQS*_125) or the full-length riboswitch prior to the termination point (*GlyQS*_213) were attached to the microscope slide through the 5’-biotinylated nascent RNA transcript^14^ (Fig. 4a and b). Upon simultaneous injection of rNTPs and U46^Cy5^-tRNA, ligand sensing was visualized co-transcriptionally in real-time until the RNAP reached the end of the 5’-Cy3-labeled DNA template strand, as indicated by the occurrence of a PIFE signal (Protein Induced Fluorescence Enhancement)^39^. For kinetic analysis, active transcription elongation complex traces were selected based on the occurrence of the PIFE signal and display of at least one U46^Cy5^-tRNA binding event either co- or post-transcriptionally.

**Fig 4.**
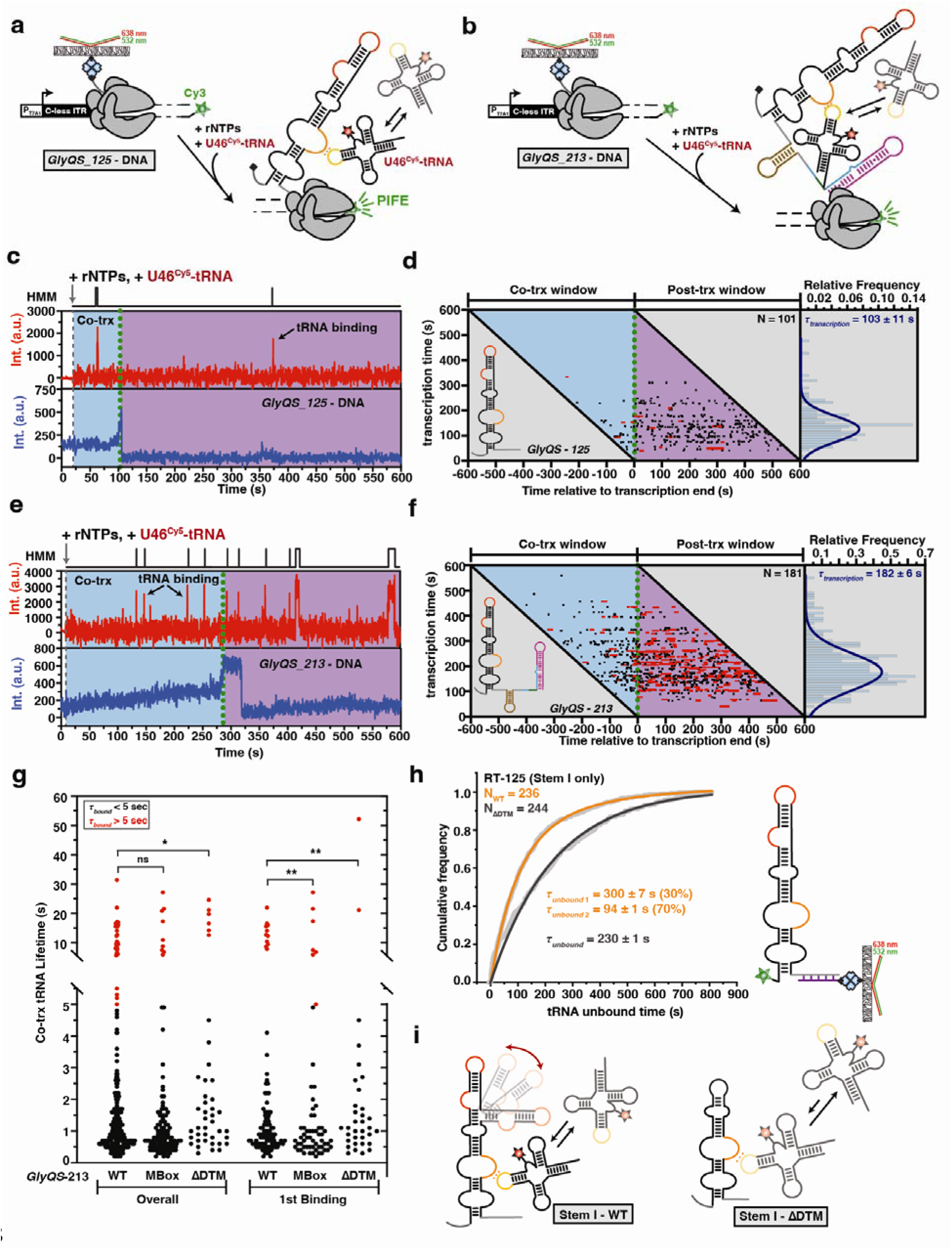
Real-time Monitoring of Ligand Binding during Transcription. **a-b**, Transcription of the DNA templates labeled with Cy3 at the 3′-end using the *E. coli* RNAP allows the formation and detection of a fluorescent Halted Complex (EC-13) attached to the microscope slide through the nascent RNA transcript. DNA templates encoding either Stem I only (**a**) or the full-length riboswitch prior to the termination point (**b**) were used in this study. Transcription is re-started upon addition of all rNTPs and addition of U46^Cy5^-tRNA in the transcription mix allows surveying co-transcriptional ligand binding in real-time. When the RNAP reaches the end of the DNA template, the occurrence of the PIFE signal (Protein Induced Fluorescence Enhancement) delimits the co-transcriptional window. **c**, Representative single molecule trajectory showing the real-time transcription of the *GlyQS*_125 DNA transcribed in the presence of 25 µM rNTPs and 25 nM U46^Cy5^-tRNA. Transcription restart is indicated by rNTPs injection, and the end of transcription is indicated by PIFE (green dotted line). Repeated co-transcriptional bindings of U46^Cy5^-tRNA are monitored upon direct excitation of the Cy5 fluorescent dye. Hidden Markov Modeling (HMM) is indicated on the top of the trace. **d**, Rastergram of U46^Cy5^-tRNA binding (black and red bars) during (cyan background) and just after transcription (magenta background) of the *GlyQS*_125 DNA template; each horizontal line represents a single transcribed molecule. The lengths and colors of horizontal bars indicate the lifetimes of individual complexes. On the x axis, t = 0 represents the time relative to the transcription end marked by occurrence of the PIFE signal. Histogram of transcription time is indicated on the right. The indicated error is the error of the gaussian fit. **e**, Representative single molecule trajectory showing the real-time transcription of the *GlyQS*_213 DNA transcribed in the presence of 25 µM rNTPs and 25 nM U46^Cy5^-tRNA. Transcription restart is indicated by rNTPs injection, and the end of transcription is indicated by PIFE (green dotted line). Repeated co-transcriptional bindings of U46^Cy5^-tRNA are monitored upon direct excitation of the Cy5 fluorescent dye. Hidden Markov Modeling (HMM) is indicated on the top of the trace. **f**, Rastergram of U46^Cy5^-tRNA binding (black and red bars) during (cyan background) and just after transcription (magenta background) of the *GlyQS*_213 DNA template; each horizontal line represents a single transcribed molecule. The lengths and colors of horizontal bars indicate the lifetimes of individual complexes. On the x axis, t = 0 represents the time relative to the transcription end marked by occurrence of the PIFE signal. Histogram of transcription time is indicated on the right. The indicated error is the error of the gaussian fit. **g**, Plot displaying the distribution of tRNA lifetimes during the transcription (co-trx) of the *GlyQS*_213 DNA templates WT, Mbox and ΔDTM. Short binding events (τ*_bound_* <5 s) are indicated in black and long-lived binding events (τ*_bound_* >5 s) are indicated in red. The total number of molecules analyzed is as follows: WT = 181; Mbox = 61; ΔDTM = 35. (****P < 0.001, ***P < 0.005, **P < 0.01, *P < 0.05). **h**, Plots displaying the cumulative unbound dwell times of U46^Cy5^-tRNA for the released transcript comprising Stem I only (RT-125) in the WT and ΔDTM context with 12.5 nM U46^Cy5^-tRNA. Unbound (τ*_unbound_)* times are indicated with the error of the fit. **i**, Secondary structural model of Stem I folding in the WT and ΔDTM context.

Consistent with our data from the stalled PEC-138 and released corresponding RT-125, PIFE-monitored real-time transcription of the *GlyQS*_125 template yielded few and short-lived U46^Cy5^-tRNA binding events both during and after transcription (Fig. 4c and d). Supporting the functionality of the on-slide transcribed *Bsu-GlyQS* T-box RNAs, the post-transcriptional U46^Cy5^-tRNA bound time (∼1 s) closely matches observations with pre-transcribed RNAs of the same length (Supplementary Fig. 5a)^25^. Given the average transcription time of 103 ± 11 sec (Fig. 4d), we expected to observe 1-2 tRNA binding events per molecule, based on the average unbound time of U46^Cy5^-tRNA^Gly^ at this concentration (Supplementary Fig. 2a). By comparison, transcription of the longer *GlyQS*_213 template took 182 ± 6 s (Fig. 4f), which correlates with the higher number of co-transcriptional U46^Cy5^-tRNA binding events―1.4 (253/181) binding events per molecule for *GlyQS*_213 compared to 0.28 (29/101) events per RT-125 molecule (Fig. 4e,f). Importantly, this longer template also displayed a substantial increase in long-lived tRNA binding events (∼11% among the total number of binding events), indicating that stable tRNA^Gly^ anchoring can occur during ongoing transcription (Fig. 4f and g). Notably, this fraction of long-lived events further increased post-transcriptionally (∼30% - Fig. 4f, Post-Trx window), suggesting an antagonism between active transcription and stabilized tRNA anchoring that is overcome after the RNAP has passed on, reminiscent of S4 protein recruitment to the 16S rRNA dynamics during 16S rRNA synthesis^38^.

To further elucidate the determinants leading to efficient co-transcriptional tRNA sensing, we performed PIFE-monitored real-time transcription of our *GlyQS*_213 templates bearing mutations that disrupt either tRNA binding to Stem I (ΔDTM) or recognition of the tRNA 3′-end (Mbox; Supplementary Fig. 4g and 5d,e). The overall co-transcriptional lifetime of U46^Cy5^-tRNA on the *GlyQS*_213-Mbox transcript was comparable to that of the WT transcript, suggesting that the tRNA can still be sensed efficiently even in the absence of a functional T-box motif. However, kinetic analysis revealed that the fraction of long-lived U46^Cy5^-tRNA:*GlyQS*_Tbox RNA complexes was substantially reduced for the Mbox mutant compared to WT, dropping to 1.2 (118/96) binding events per molecule, supporting an essential role of the T-box motif in stable tRNA^Gly^ anchoring during transcription (Supplementary Fig. 5d). Consistent with this conclusion, observation of only few (11/270, or 4%, relative to the WT’s ∼30%) long tRNA bound times after transcription completion (Post-Trx window) of the *GlyQS*_213-Mbox transcript similarly demonstrated that the T-box motif contributes substantially to tRNA anchoring under thermodynamic equilibrium conditions (Supplementary Fig. 5b).

**Fig 5.**
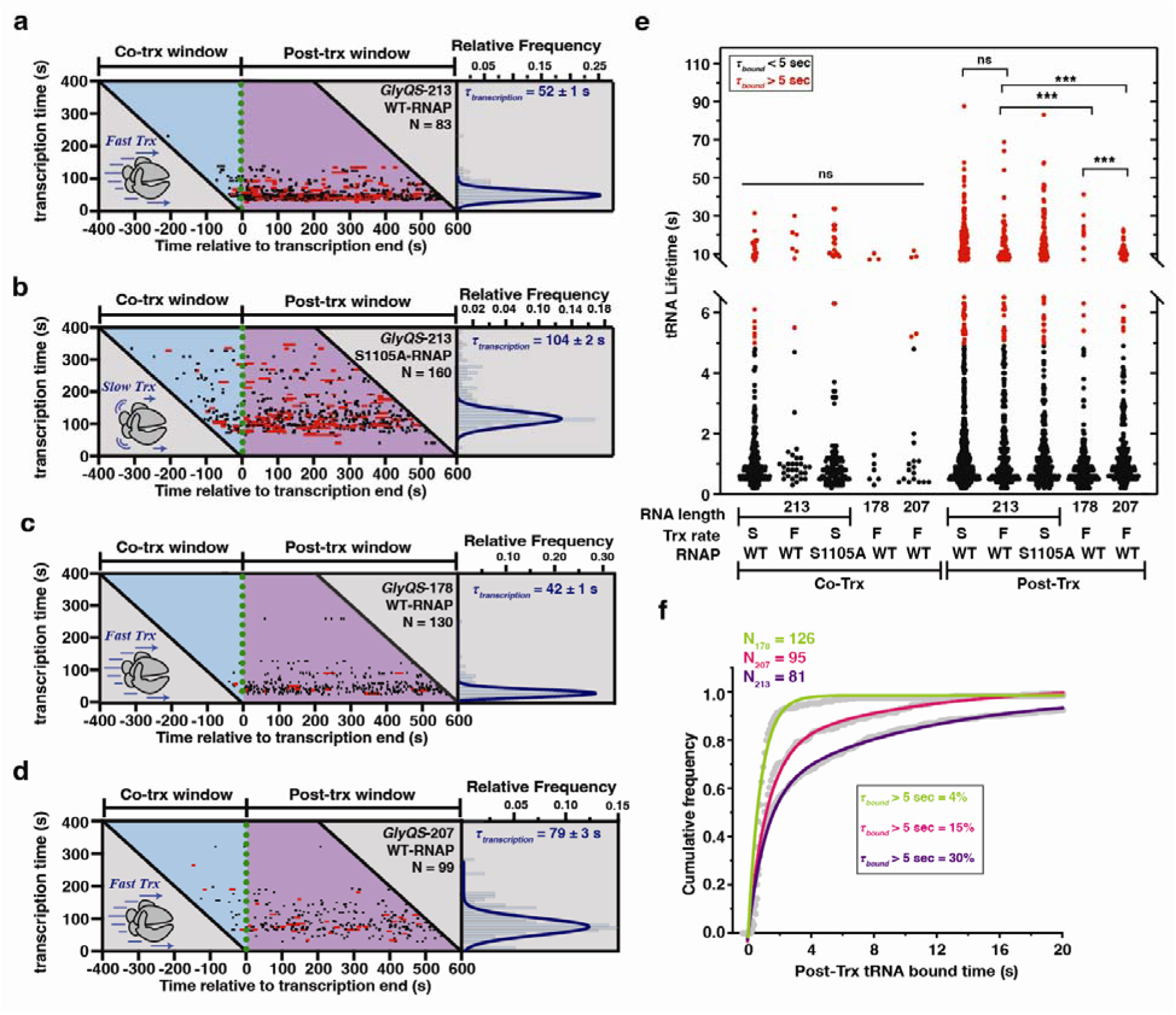
RNAP Pausing Provides a Kinetic Window Allowing Efficient Ligand sensing. **a-b**, Rastergrams of U46^Cy5^-tRNA binding (black and red bars) during (cyan background) and just after transcription (magenta background) using 250 µM rNTPs for the transcription of the *GlyQS*_213 DNA template with WT (**a**) and slow RNAP (**b**); each horizontal line represents a single transcribed molecule. The lengths and colors of horizontal bars indicate the lifetimes of individual complexes. On the x axis, t = 0 represents the time relative to the transcription end marked by occurrence of the PIFE signal. Histogram of transcription time is indicated on the right. The indicated error is the error of the gaussian fit. **c-d**, Rastergrams of U46^Cy5^-tRNA binding (black and red bars) during (cyan background) and just after transcription (magenta background) using 250 µM rNTPs for the transcription of the *GlyQS*_178 (**c**) and *GlyQS*_207 (**d**) DNA templates with WT RNAP; each horizontal line represents a single transcribed molecule. The lengths and colors of horizontal bars indicate the lifetimes of individual complexes. On the x axis, t = 0 represents the time relative to the transcription end marked by occurrence of the PIFE signal. Histogram of transcription time is indicated on the right. The indicated error is the error of the gaussian fit. **(e)** Plot displaying the distribution of tRNA lifetimes during (cyan) and after the transcription (magenta) of the *GlyQS*_213, *GlyQS*_178 and *GlyQS*_207 DNA templates using slow (S) and fast (F) transcription rates. Short binding events (τ*_bound_* <5 s) are indicated in black and long-lived binding events (τ*_bound_* >5 s) are indicated in red. (****P < 0.001, ***P < 0.005, **P < 0.01, *P < 0.05). **f**, Plots displaying the cumulative bound dwell times of U46^Cy5^-tRNA after transcription completion of the *GlyQS*_213, *GlyQS*_178 and *GlyQS*_207 DNA templates. The proportions of the long-lived bound times (τ*_bound_* >5 s) relative to the overall bound time are indicated.

In contrast, the *GlyQS*_213-ΔDTM transcript produced few and dominantly short-lived tRNA^Gly^ binding events (Fig. 4g and Supplementary Fig. 5c), with only 0.47 (41/87) long-lived binding events per molecule remaining, suggesting that early co-transcriptional recruitment via Stem I is critical for effective downstream ligand anchoring during transcription elongation; that is, anchoring engages both the tRNA elbow and 3′-end interaction points.

To further test this model, we examined the dwell times of the first U46^Cy5^-tRNA binding interaction during active transcription (Fig. 4g, right panel). For the WT, we observed both short-and long-lived binding events, consistent with prior models in which initial tRNA association occurs independently of anchoring^24–26^, which is established only after AT transcription and formation of Stems I and III, and most efficiently when both Stem I and the T-box motif are displayed outside the RNAP exit channel (Fig. 4g). In contrast, both the Mbox and ΔDTM mutants exhibited a pronounced reduction in long-lived binding events, with the effect being strongest for the ΔDTM mutant (Fig. 4g). Together, these results support a model in which early initial ligand recruitment via Stem I is a prerequisite for subsequent, stable tRNA anchoring during transcription.

Kinetic analysis of U46^Cy5^-tRNA binding to the co-transcriptionally folded, RNAP-free RT-125 RNA revealed that Stem I alone senses tRNA^Gly^ via two kinetic regimes (Fig. 4h). One component is a comparably short unbound time (∼94 s), while the other is substantially longer (∼300 s), likely to represent a conformation with tRNA binding events too sparse to be functional under cellular conditions. Notably, the RT-125-ΔDTM transcript displayed only the slower kinetic regime (230 s unbound time; Fig. 4h and i), underscoring the importance of Stem I folding for early tRNA^Gly^ recruitment during transcription, as previously suggested^24,25^.

When PIFE-monitored real-time transcription was performed with the tRNA 3′ end blocked in Gly-tRNA-3′-Cy5, very few co-transcriptional binding events were observed, further supporting the notion that discrimination of the aminoacylation state is actively performed during transcription (Supplementary Fig. 5d). Together, our real-time analysis of co-transcriptional U46^Cy5^-tRNA binding uncovers a hierarchical mechanism of ligand recognition, in which early recruitment to a properly folded Stem I (sensing) enables efficient downstream tRNA^Gly^ capture (anchoring) to the T-box motif.

### Transcription rate and pausing set a kinetic checkpoint for efficient tRNA sensing

Co-transcriptional processes such as RNA folding, protein recruitment and ligand recognition are tightly and rapidly coordinated to precisely tune gene regulatory responses to ever-changing environmental cues^2,40^. In riboswitch systems, the transcription rate is an additional key determinant of ligand recognition and the resulting gene regulation, by enforcing a kinetic window during which sensing has to occur^3,7,11,13,14^.

To assess the impact of the transcription rate on tRNA^Gly^ sensing, we increased the concentration of rNTPs from 25 µM to 250 µM, accelerating T-box RNA elongation during PIFE-monitored real-time transcription of the *GlyQS*_213 template (Fig. 5a). This 10-fold increase led to an ∼4-fold rise in transcription rate^41^, taking only 52 ± 1 s of mean transcription time (Fig. 5a) compared to 182 ± 6 s at 25 µM rNTPs (Fig. 4f). This shortened time markedly reduced the number of co-transcriptional U46^Cy5^-tRNA binding events, consistent with the narrower kinetic window available for ligand sensing. Conversely, an RNAP S1105A mutant, which elongates 2-fold more slowly at 250 µM rNTPs than WT RNAP, partially restored U46^Cy5^-tRNA sensing opportunity due to a longer mean transcription time (104 ± 2 s, Fig. 5b). We note that increased transcription rates generally were accompanied by a narrowing of the transcription time distribution across single transcription complexes (compare Figs. 4f and 5a-d). Together, these observations indicate that rNTP availability―and the resulting transcription rate as a sensor of cellular energy status―sets the kinetic window for co-transcriptional tRNA^Gly^ recognition.

Strikingly, despite these rate-dependent changes in overall sensing opportunity, the fraction of long-lived U46^Cy5^-tRNA:*GlyQS* T-box interactions remained essentially constant (∼15%) across all transcription rates tested (Fig. 5e and Supplementary Fig. 6c,d). This finding suggests that, although fast transcription narrows the sensing window, stable tRNA^Gly^ anchoring still occurs with unchanged probability under these more physiological elongation conditions, where intracellular rNTP concentrations are estimated to be in the low-millimolar range^42^. Consistent with this observation, transcription termination assays conducted under both fast and slow transcription regimes yielded similar termination and tRNA^Gly^ binding efficiencies (Supplementary Fig. 6a,b).

Next, to examine how transcriptional pausing shapes co-transcriptional riboswitch folding and ligand sensing, we performed PIFE-monitored real-time transcription of DNA templates truncated at the 178 and 207 pause sites (Fig. 5c and d). Consistent with our prior assays using similarly truncated, co-transcriptionally folded RTs (Supplementary Fig. 2e,f), nascent RNAs produced from these templates displayed mostly sparse, short-lived U46^Cy5^-tRNA binding events during transcription (Fig. 5e). Closer analysis of the bound times revealed that tRNA^Gly^ binding slightly increases both co- and post-transcriptionally as the transcript extends from Stem I (178 pause) to completion of the AT/T switching region (207 pause, Fig. 5e).

Strikingly, the full-length *GlyQS*_213 real-time transcript, which includes a complete T sequence, displayed substantially more long-lived U46^Cy5^-tRNA binding events than the only slightly shorter *GlyQS*_207 transcript (Supplementary Fig. 6e). This dependence on the final nucleotides is consistent with an off-pathway fold involving nucleotides 202-206 within the nascent T hairpin that compete with formation of the productive AT hairpin (nucleotides 178-182) required to engage the tRNA 3′ end, until additional T-hairpin sequence is transcribed. Consistent with this model, dwell-time analysis revealed a substantially higher fraction of co-transcriptional U46^Cy5^-tRNA:T-box complexes for *GlyQS*_213 than for *GlyQS*_207 (Fig. 5f), delineating the transcriptional window available for stable tRNA anchoring. Accordingly, in a purely post-transcriptional binding context, the fraction of long-lived complexes was substantially reduced from 41% for *GlyQS*_213 to 24% for *GlyQS*_207 (Supplementary Fig. 6e).

Together, these findings demonstrate that tRNA^Gly^ recruitment and full (stable) anchoring occur within a narrow transcriptional window—a kinetic checkpoint governed by RNAP elongation rate and pause-site positioning—and depend on timely formation of the thermodynamically less favored AT hairpin immediately before the termination point.

### Real-time correlation of ligand binding with transcription unveils the readthrough mechanism

Ligand binding is central to riboswitch-mediated gene regulation in response to cellular cues^43^. To directly link tRNA^Gly^ binding to downstream transcriptional output, we aimed to extend our real-time transcription assay to monitoring regulation at the single-molecule level (Fig. 6). Because both intrinsic termination and readthrough followed by run-off result in RNAP and DNA dissociation from the surface-tethered transcript—rendering these outcomes indistinguishable in our standard format—we implemented a double-PIFE strategy^44^ to resolve them. We designed a doubly labeled DNA template (*GlyQS_207^Cy3^_247^Cy3^*) bearing Cy3 fluorophores at two positions: an internal label at position 207 to report RNAP passage through the final pause site, and a second label at the downstream template end to report transcription beyond the poly-U tract and subsequent run-off (Fig. 6a,b).

**Fig 6.**
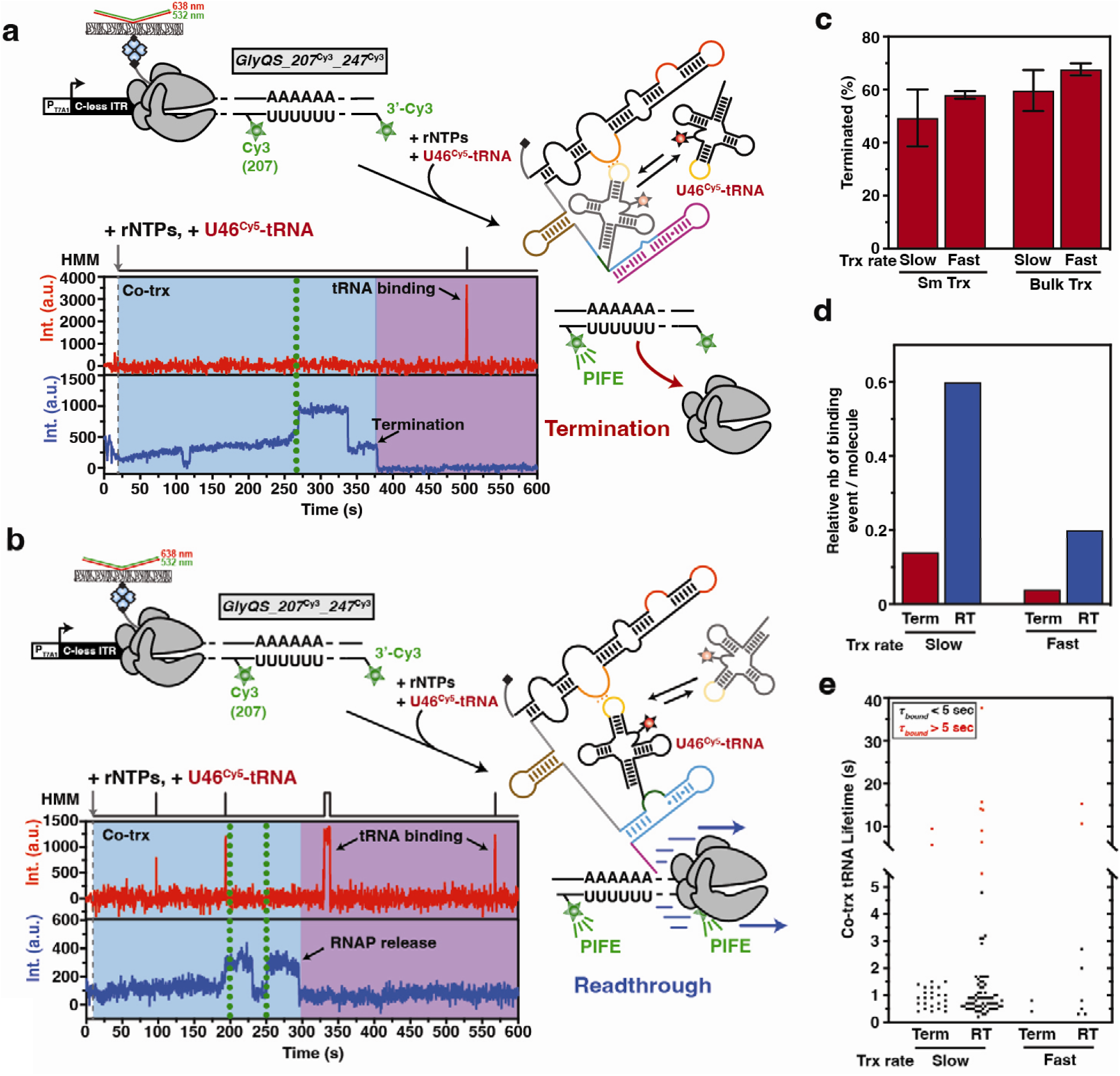
Monitoring Transcription Regulation at the Single-molecule Level. **a-b**, Transcription of the DNA template labeled with two Cy3 fluorophores at the 207 position and downstream of the Poly-U sequence allows for the detection of transcription termination (a) and transcriptional readthrough (b) at the single molecule level. Transcription is re-started upon addition of all rNTPs and addition of U46^Cy5^-tRNA in the transcription mix allows surveying co-transcriptional ligand binding in real-time. **c**, Transcription termination efficiencies determined during real-time transcription (Sm Trx) and during bulk transcription performed *in vitro* (Bulk Trx). Transcriptions were performed using 25 µM (Slow) or 250 µM (Fast) rNTPs with 25 nM U46^Cy5^-tRNA. The total number of molecules analyzed is as follows: Terminated_slow_ = 176; Readthrough_slow_ = 119; Terminated_fast_ = 50; Readthrough_fast_ = 37. **d**, Relative number of U46^Cy5^-tRNA binding events detected per molecule transcribed in real-time in the context of terminated (Red) and readthrough (Blue) transcripts. Transcriptions were performed using 25 µM (Slow) or 250 µM (Fast) rNTPs with 25 nM U46^Cy5^-tRNA. **e**, Plot displaying the distribution of tRNA lifetimes during the transcription of the *GlyQS*_207^Cy3^_247^Cy3^ DNA template during slow and fast transcription rates. Short binding events (τ*_bound_* <5 s) are indicated in black and long-lived binding events (τ*_bound_* >5 s) are indicated in red. The total number of molecules analyzed is as follows: Terminated_slow_ = 176; Readthrough_slow_ = 119; Terminated_fast_ = 50; Readthrough_fast_ = 37.

Real-time transcription of this template―initiated by simultaneous rNTP and U46^Cy5^-tRNA injection―yielded two classes of single molecule fluorescence traces. Termination events were characterized by a single PIFE signal originating from position 207, followed by loss of the signal upon RNAP and (likely) DNA dissociation at the termination point poly-U (Fig. 6a). In contrast, readthrough events engendered two distinct PIFE signals, indicating transcription through both the pause site 207 and beyond the termination point to the end of the template (Fig. 6b). This design enables direct visualization of transcription termination regulation at single-molecule resolution.

To validate that this DNA template produces functional *Bsu-GlyQS* T-box riboswitches, we compared termination efficiency on-slide with that measured in bulk transcription assays analyzed by gel electrophoresis. Real-time transcription of the *GlyQS_207^Cy3^_247^Cy3^* template resulted in 49 ± 11% and 58 ± 1% termination at low (25 µM) and high (250 µM) rNTP concentrations―i.e., slow and fast transcription rates), respectively. The close match with bulk results (Fig. 6c) confirms the suitability of our double-PIFE assay in assessing gene regulation outcomes.

Next, we asked how initial tRNA^Gly^ recruitment and subsequent anchoring relate to transcriptional fate. Readthrough traces exhibited ∼4-fold more U46^Cy5^-tRNA binding events than prematurely terminated traces (Fig. 6d), consistent with ligand engagement promoting antitermination. This relationship held at both slow and fast transcription rates: Although fast transcription (high rNTP) reduced the total number of binding events, the ∼4-fold enrichment in readthrough versus termination was preserved (Fig. 6d). Notably, U46^Cy5^-tRNA:T-box complexes associated with terminated transcripts were uniformly short-lived (<1 s; Fig. 6e), whereas long-lived (>5 s) complexes were observed only in readthrough events—most prominently under fast transcription (Fig. 6d)—indicating that stable tRNA^Gly^ anchoring is required for productive antitermination under near-physiological conditions.

Together with the data in Figures 4 and 5, these results support a model in which early tRNA^Gly^ recruitment to Stem I, followed by AT-hairpin-mediated 3′-end anchoring, actively promotes transcriptional readthrough, directly illuminating how this riboswitch executes gene regulation under cellular conditions.

## Discussion

Riboswitches are co-transcriptional molecular switches that modulate gene expression—most commonly by controlling transcription termination or translation initiation—through recognition of cognate ligands^2^. Here, we directly couple ligand binding dynamics to transcriptional outcome at the single-molecule level, providing a real-time mechanistic view of how the *Bsu-GlyQS* T-box riboswitch senses uncharged tRNA^Gly^ and drives antitermination. These observations—shaped by strategically positioned pause sites—reveal how the riboswitch exploits directional, hierarchical 5’-to-3′ directional, folding to create a defined kinetic window for ligand engagement and efficient regulation of downstream gene expression (Fig. 7).

**Fig. 7.**
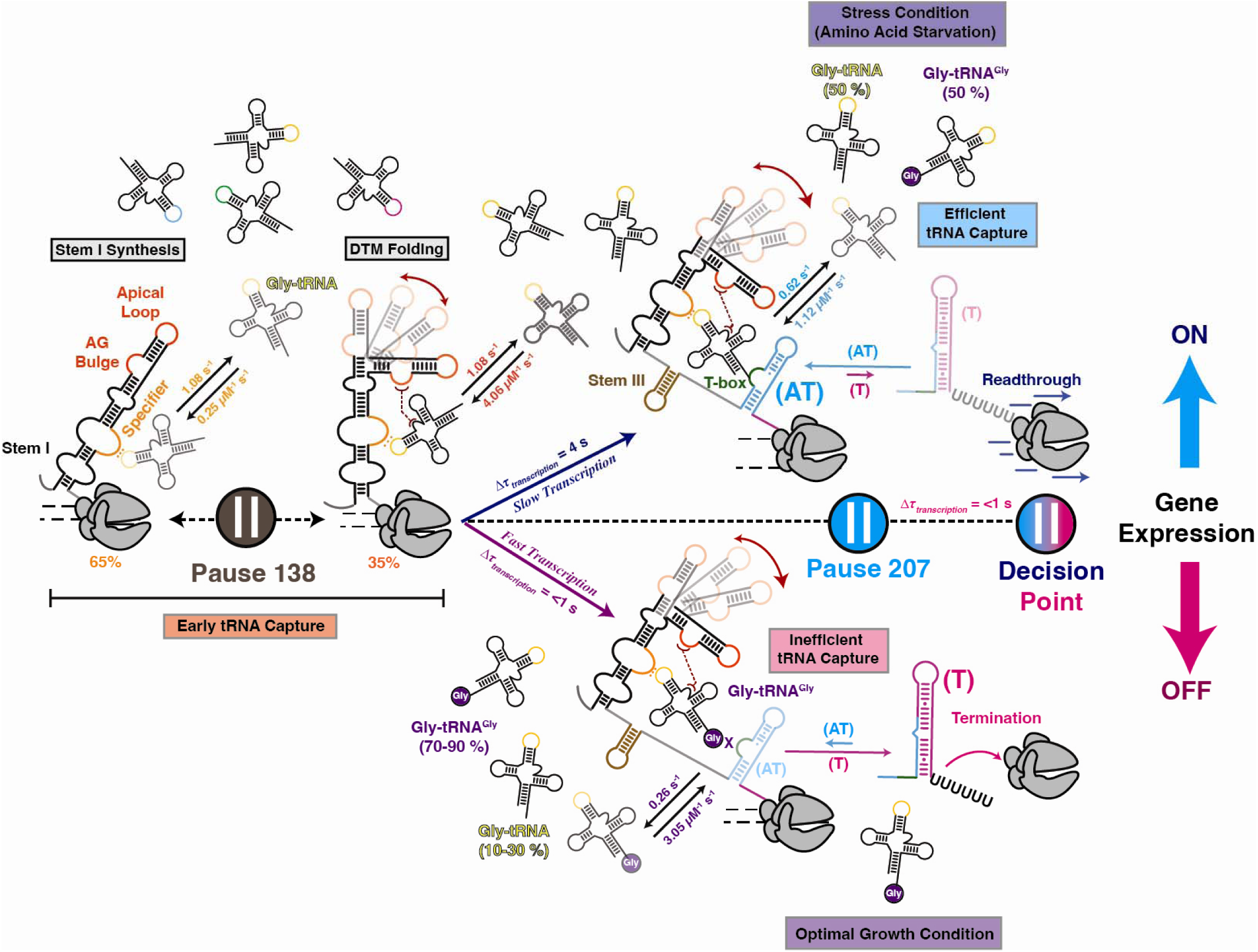
Co-transcriptional Model of Gly-tRNA Sensing to the *Bsu-GlyQS* T-box Riboswitch. Recruitment of tRNA^Gly^ begins once Stem-I, with the DTM and the Specifier sequence, emerges from the RNA polymerase (RNAP). The first transcriptional pause site (138) allows more time for proper folding of the DTM and early tRNA^Gly^ engagement. Stem-I binds both the charged and uncharged tRNAs with similar k_on_ and k ^31^. Under stress condition (top panel), transcription is slowed down (*τ_transcription_*∼4 s), favoring the anchoring of uncharged tRNA^Gly^. Moreover, a longer-lasting pause 207 prior to the decision point allows additional time for AT folding and efficient anchoring of the tRNA^Gly^ 3′-end. In the absence of stress (bottom panel), transcription progress is fast (τ*_transcription_* <1 s), directly competing with ligand anchoring (bound time ∼3 s). In addition, under this cellular condition the majority of tRNA^Gly^ molecules will be charged, therefore creating an additional competition barrier that suppresses uncharged tRNA^Gly^ anchoring, thereby down-regulate the expression of downstream genes.

The *Bsu-GlyQS* T-box riboswitch follows a hierarchical folding program that enables efficient recruitment and stable anchoring of uncharged tRNA^Gly^, thereby biasing transcription toward readthrough rather than termination (Fig. 1). Our results support a dual mechanism in which maturation of the riboswitch scaffold is tightly coupled to dynamic tRNA recognition, with Stem I and the downstream T-box motif playing distinct, sequential roles. Stem I is essential for establishing the global architecture of the aptamer and for initiating a two-stage, co-transcriptional binding pathway previously inferred for translational T-box systems^24^: Early, transient interactions mediated by Stem I—captured here as U46^Cy5^-tRNA binding events—pre-organize the complex and promote the structural register required for subsequent 3′-end anchoring later in transcription (Fig. 7). Within Stem I, the DTM strengthens specifier-anticodon recognition and stabilizes adoption of the functional conformation, endowing Stem I with a chaperone-like role. In this respect, Stem I parallels the bacterial RNA-binding protein Hfq, which promotes productive RNA-RNA interactions by stabilizing and coordinating RNA conformational transitions^45,46^. By stabilizing early tRNA engagement (Figs. 2 and 4), Stem I primes the riboswitch for downstream T-box-mediated 3′end anchoring during a key RNAP pausing event, enabling a seamless transition to the fully bound, antiterminating state (Fig. 7).

Our kinetic analyses uncover tight coupling between riboswitch folding and RNAP-mediated pausing, which extends the time available for AT-region rearrangements and for tRNA sensing. Formation of the AT hairpin exposes the T-box motif and enables recognition of the tRNA 3′ end—an essential step in committing the complex to antitermination and transcriptional readthrough. Consistent with this model, mutagenesis establishes that both Stem I and the T-box motif are required to support stable tRNA binding, underscoring the functional importance of hierarchical folding in gene regulation^25^. More broadly, the ordered assembly pathway we observe mirrors principles of ribosomal RNA folding and RNP complex formation^4^, where temporally staged interactions enforce specificity and robustness. In this view, RNAP pausing operates as a kinetic checkpoint that coordinates successive structural transitions, analogous to promoter-proximal pausing in developmental gene control^47^. By gating progression until upstream steps are completed, such checkpoints help preserve the fidelity, timing, and efficiency of gene-expression decisions.

T-box riboswitches are highly adaptive regulators that couple amino acid availability to gene expression by sensing changes in the abundance of uncharged tRNA. Our results show that conformational plasticity during elongation enables the riboswitch to tune transcriptional output as a function of both RNAP elongation rate and tRNA availability (Figs. 4,5). In particular, competition between charged and uncharged tRNA species provides a dynamic input that can shift riboswitch behavior and thereby shape the transcriptional response to nutrient stress (Fig. 7). In cells, metabolic state strongly influences the fraction of charged versus uncharged tRNA^48^, a key determinant of T-box function. Under nutrient-starved conditions, ∼70–90% of tRNAs are typically aminoacylated, leaving only a small pool of uncharged Gly-tRNA (estimated at ∼0.2–1 μM)^49–52^. In this regime, abundant charged Gly-tRNA can occupy early binding contacts yet fail to support the critical 3′-NCCA:T-box anchoring interaction required for AT stabilization and terminator destabilization, effectively biasing the switch toward termination (Fig. 7). During glycine starvation, by contrast, rising levels of uncharged tRNA^Gly^ promote 3′-end anchoring and antitermination, upregulating genes involved in amino acid biosynthesis, transport, and tRNA charging. Given glycine’s roles in protein synthesis, one-carbon metabolism, and broader intermediary metabolism^53^, fluctuations in its availability position the T-box riboswitch as an integrative sensor of cellular nutritional state.

In summary, the *Bsu-GlyQS* T-box riboswitch illustrates how hierarchical folding, strategically positioned transcriptional pauses, and ligand availability converge to drive efficient, responsive control of gene expression. The mechanistic parallels to other RNA-guided pathways, including Hfq-facilitated sRNA–mRNA pairing, underscore broader principles of RNA-based regulation. Notably, the T-box RNA itself exhibits chaperone-like activity, promoting productive hybridization between the tRNA 3′ end and the T-box motif. By directly linking co-transcriptional tRNA binding kinetics to transcriptional fate in real-time, the current study establishes a first quantitative framework and experimental blueprint for atomic-resolution dissection of these transitions, for testing how shared design principles extend across riboswitch and RNA regulatory networks, and for ultimately informing strategies to therapeutically perturb T-box-controlled pathways in bacterial pathogens.

## Materials and Methods

### DNA templates and Oligonucleotides

A 349-nucleotide DNA template including the *GlyQS-*TboxT-box riboswitch from *B. subtilis* under the control of the T7A1 promoter was cloned into pUC-GW plasmid (Azenta Genewiz). Primers were purchased from Integrated DNA technologies, Inc. (IDT) with HPLC purification when containing modifications. Transcription templates for *in vitro* transcription were generated by PCR using the “T7A1-PCR” forward oligonucleotide and the according reverse oligonucleotides. For mutant DNA templates, the mutation was inserted in two PCR steps using overlapping oligonucleotides containing the corresponding mutation.

Fluorescent DNA templates for transcription were generated by PCR using Phusion high fidelity polymerase (NEB). Following PCR, fluorescently labeled DNA templates were separated on 1% agarose, column purified and eluted in 40 µL of water.

For the DNA template containing two Cy3 fluorophores, reverse primer (see key resources table) containing an internal amino-alkyl modified nucleotide (IDT; see Supplementary Table 2) was fluorescently labeled with Cy3-NHS dye (Lumiprobe) as follows: 0.5 mg of Cy3 dye was dissolved in 33 µL of DMSO and added to a reaction mixture containing 5 nmol modified primer adjusted to 100 µL final volume with 100 mM sodium bicarbonate pH 8.5. Reactions were incubated overnight at room temperature and then free dye was removed using a Chroma TE-10 spin column (Takara Bio) followed by clean-up on a Monarch DNA column using the oligonucleotide clean-up protocol.

Oligonucleotides used in this study are listed in Supplementary Table 1.

### *In vitro* transcription assays

Halted complexes (EC-13) were prepared in transcription buffer (20 mM Tris-HCl, pH 8.0, 20 mM NaCl, 20 mM MgCl_2_, 14 mM 2-mercaptoethanol, 0.1 mM EDTA) containing 25 µM ATP/UTP mix, 50 nM α^32^P-GTP (3000 Ci/mmol), 10 µM ApC dinucleotide primer (Trilink), and 50 nM DNA template. *E. coli* RNAP holoenzyme (New England Biolabs) was added to 100 nM, and the mixture was incubated for 10 min at 37 °C. The sample was passed through G50 column to remove any free nucleotides. To complete the transcription reaction all four rNTPs were added concomitantly with heparin (450 µg/mL) to prevent the re-initiation of transcription. The mixture was incubated at 37 °C, and reaction aliquots were quenched at the desired times into an equal volume of loading buffer (95% formamide, 1 mM EDTA, 0.1% SDS, 0.2% bromophenol blue, 0.2% xylene cyanol). Reaction aliquots were denatured before loading onto a denaturing 8 M urea, 6% polyacrylamide sequencing gel. The gel was dried and exposed to a phosphor screen (typically overnight), which was then scanned on a Typhoon Phosphor Imager (GE Healthcare).

### Transcription data analysis

To determine the RT_50_ for regulation of termination by Gly-tRNA, the relative intensity of the full-length product was divided by the total amount of RNA transcript (full-length + terminated product) for each Gly-tRNA concentration. The resulted percentages of readthrough were plotted against the Gly-tRNA concentration using equation: Y=Y_0_ + ((M1×X)/(T_50_+X)), where Y_0_ is the value of Y at X_min_ (no ligand condition) and M1 = Y_max_ - Y_min_. Error bars in transcription quantification represent the standard deviation of the mean from independent replicates.

### Aminoacylation of Gly-tRNA and 3′-end labeling

Aminoacylation of Cy5-labeled Gly-tRNA was achieved using E. coli S100 extracts (tRNA probes, College Station, TX, USA) in a 25 μl reaction volume following a protocol previously described^54^. Briefly, ∼2 μM of Cy5-tRNAGly was charged with 3 μl of E. coli S100 extract in a 50 μl reaction containing 100 mM HEPES-KOH, pH 7.6, 200 μM glycine, 10 mM ATP, 1 mM dithiothreitol, 20 mM KCl, 20 mM MgCl_2_, and incubated at 37 °C for 30 min. The reaction was stopped by adding 3 M sodium acetate, pH 5.2, to a final concentration of 100 mM in 100 μl volume. Aminoacylated tRNA was purified by performing phenol–chloroform extraction to remove the proteins. The concentration of the Gly-tRNA^Gly^ was measured using a nanodrop 2000 and multiple aliquots of it were made and stored at −80 °C. An acid-urea PAGE was used for checking the aminoacylation efficiency of the reaction. Under our reaction conditions, the aminoacylation efficiency was close to ∼90%^25^. For the single-molecule binding experiments, Cy5-labeled Gly-tRNA^Gly^ was diluted to the required concentration in standard buffer.

For 3′-end labeling of the Gly-tRNA with Cy5 fluorophore, we employed RNA ligation with pCp-Cy5 (Jena Bioscience). Briefly, in a 30 µL reaction volume, 3 pmol of unlabeled Gly-tRNA were incubated with 30 pmol pCp-Cy5, 1 mM ATP, 10% DMSO (v/v) and 1 U of T4 RNA ligase I (NEB) overnight at 16 °C in the reaction buffer provided by the supplier. Labeling reaction was run on a 20% denaturing urea PAGE and gel-excised to purify the labeled from the unlabeled product followed by ethanol precipitation.

### Preparation of fluorescently labeled nascent transcripts

*In vitro* transcription reactions were performed in two steps to allow the specific incorporation of Cy3 at the 5’-end of the RNA sequence. Transcription reactions were performed in the same transcription buffer described above. Transcription reactions were initiated by adding 100 μM 5’-Cy3-ApC and 25 μM ATP/UTP/GTP nucleotides at 37 °C for 10 min, thus yielding a fluorescent halted complex. The sample was next passed through a G50 column to remove any free nucleotides, and the transcription was resumed upon addition of all four rNTPs at the indicated concentration and heparin (450 μg/mL) to prevent the re-initiation of transcription. The resulting nascent transcript was hybridized to the 5’-biotinylated CP (Anchor Bio oligonucleotide) complementary to the 3′-end capture sequence, allowing immobilization of the complex to the microscope slide. The CP was mixed to a ratio of 10:1 with RNA transcript and added for 5 min before flowing the whole complex to the microscope slide. In the case of PEC transcription, the DNA templates contain a biotin at the 5’-end of the template DNA strand. Streptavidin was mixed to a ratio of 5:1 with the DNA template for 5 min prior to starting the transcription reaction.

For real-time transcription assays, transcription reactions were initiated by adding 100 μM 5’-Biotin-ApC (Horizon Discovery) and 25 μM ATP/UTP/GTP nucleotides at 37 °C for 10 min, thus yielding a fluorescent halted complex. The sample was next passed through a G50 column and directly flowed on microscope slide pre-incubated with streptavidin.

### Single molecule fluorescence microcopy experiments

All single-molecule fluorescence microscopy experiments were performed using an Oxford Nanoimager (ONI) S with a 100x 1.4 NA oil immersion super apochromatic objective, 405, 473, 532, 640 nm lasers with appropriate dual-band emission filters, and a Hamamatsu sCMOS Orca flash 4 V3 camera, run in TIRF mode. All movies were collected at 100 ms time resolution. PEG-passivated, biotin-modified slides with a chamber or a microfluidic channel containing inlet and outlet ports for buffer exchange were assembled as described in previous works^55^. The surface of the microfluidic channel was coated with streptavidin (0.2 mg/mL) for 10 to 15 min prior to flowing the nascent transcripts hybridized to the CP. In the case of fluorescent PECs, the complexes were directly injected into the channel surface using the biotin-streptavidin roadblock for immobilization.

For all experiments, the nascent RNA transcripts were diluted in in imaging buffer (40 mM Tris-HCl, pH 8.0, 330 mM KCl, 5 mM MgCl_2_, 0.1 mM EDTA, 0.1 mM dithiothreitol, 1 mg/mL bovine serum albumin). U46^Cy5^-tRNA (6 nM for immobilized complexes and 12 nM for real-time transcriptions) was injected into the channel in the corresponding imaging buffer along with an enzymatic oxygen scavenging system consisting of 44 mM glucose, 165 U/mL glucose oxidase from *Aspergillus niger*, 2,170 U/mL catalase from *Corynebacterium glutamicum*, and 5 mM Trolox to extend the lifetime of the fluorophores and to prevent photo-blinking of the dyes. In the case of aminoacylated U46^Cy5^-tRNA and 3′-Cy5-tRNA, 12.5 nM and 25 nM were used for immobilized complexes and real-time transcription respectively. The raw movies were collected for 10-15 min with direct green (532 nm) and red (638 nm) laser excitation.

### Single molecule data acquisition and analysis

Locations of molecules and fluorophore intensity traces for each molecule were extracted from raw movie files using custom-built MATLAB codes. Traces were manually selected for further analysis using the following criteria: single-step photobleaching of Cy3, ≥ 2 spikes of Cy5 fluorescence of more than 2-fold the background intensity. Traces showing binding events were idealized using a two-states (bound and unbound) model using a segmental *k*-means algorithm in QuB^48^. From the idealized traces, dwell times of U46^Cy5^-tRNA in the bound (τ_bound_) and the unbound (τ_unbound_) states were obtained. Cumulative plots of bound and unbound dwell-time distributions were plotted and fitted in Origin lab with single-exponential or double-exponential functions to obtain the lifetimes in the bound and unbound states. The dissociation rate constants (*k*_off_) were calculated as the inverse of the τ*_bound_*, whereas the association rate constants (*k*_on_) were calculated by dividing the inverse of the τ_unbound_ by the concentration of U46^Cy5^-tRNA used during the data collection.

tRNA binding traces that showed PIFE were plotted as 2D plots of horizontal vectors corresponding to each time trace, color coded for bound times and sorted by their time for the PIFE effect on the vertical dimension using an ad-hoc Python script.

## SUPPLEMENTAL INFORMATION

Supplemental information can be found online at…

## ACKNOWLEDGMENTS

We thank Yun Zhang for help with the Python code for real-time transcription analysis. This work was supported by NIH grants R01 GM118524 and MIRA R35 GM131922 to NGW.

## AUTHOR CONTRIBUTIONS

Conceptualization, A.C. J.C-V and N.G.W.; Methodology, A.C.; Investigation, A.C. Writing - Original Draft, A.C.; Writing - Review & Editing, A.C., J.C-V., N.G.W.; Supervision, N.G.W; Funding Acquisition, N.G.W.

## DECLARATION OF INTERESTS

The authors declare no competing interests.

## Notes

### Competing Interest Statement

The authors have declared no competing interest.

