## Supplemental Information for "Directional co-transcriptional folding and pausing create kinetic checkpoints for riboswitch-controlled gene expression"

SUPPLEMENTARY FIGURES

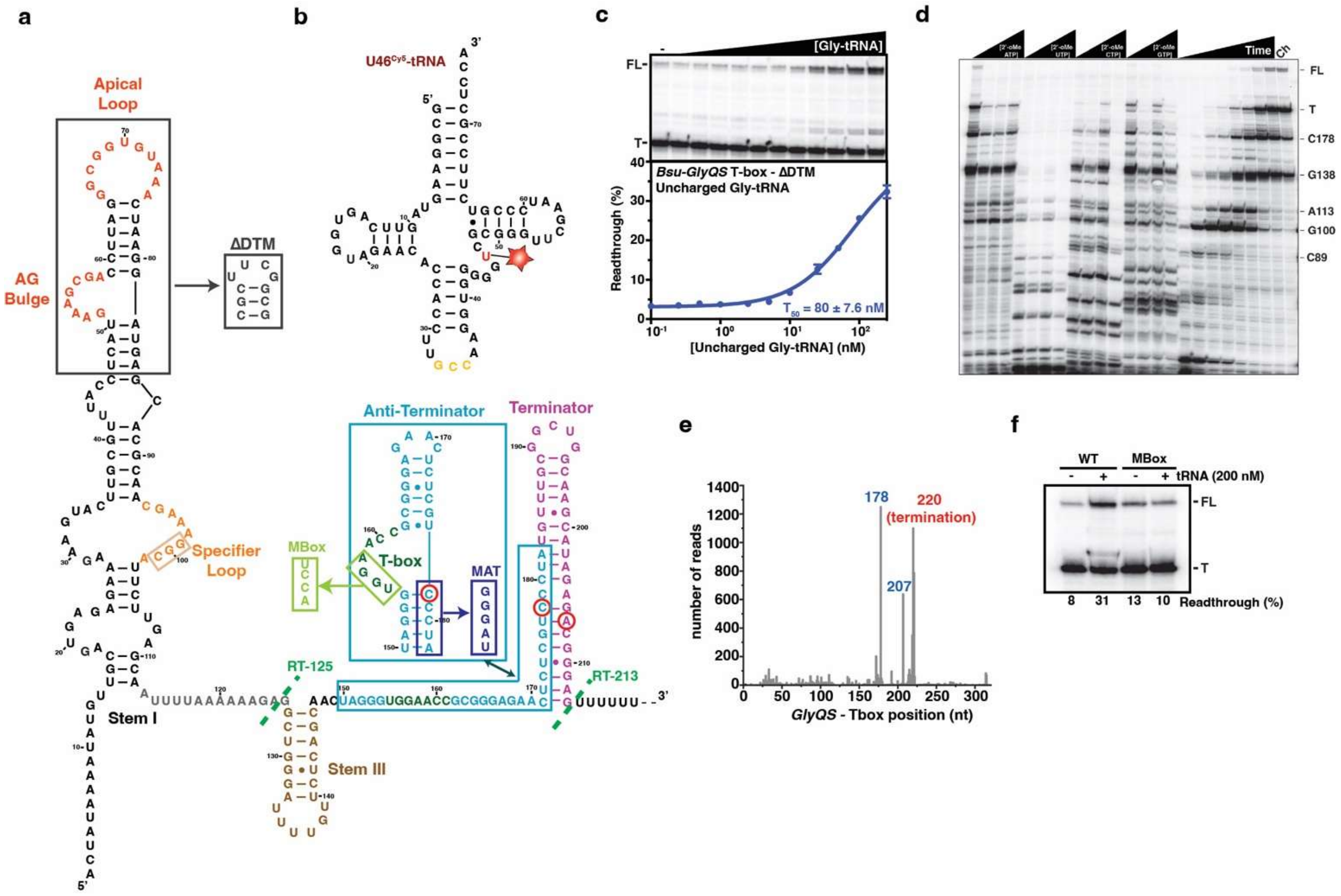


### Figure S1. Structure and regulation mediated by the *Bsu-GlyQS* T-Box riboswitch

**(a)** The entire riboswitch sequence is shown with the Stem I (black), DTM AG bulge and Apical loop (red), Specifier loop (orange), Stem III (brown), Anti-terminator (cyan), T-box motif (dark green) and the Terminator hairpin (magenta). RNAP pauses at position 178 and 207 are highlighted with a red circle. Deletion of the DTM (ΔDTM) is indicated with a grey rectangle. Mutations destabilizing the anti-terminator (MAT) are indicated in purple and mutations of the T-box motif (MBox) are indicated in light green.

**(b)** Secondary structure of the U46$^{Cy5}$-tRNA. Position of the Cy5 fluorophore is indicated in red with a star and the anti-codon is depicted in yellow.

**(c)** Representative denaturing gel (top) and plot of the fraction of transcription readthrough versus the concentration of Gly-tRNA (bottom) in the context of the ΔDTM construct. Transcription reactions were performed using 10 μM rNTPs. The $RT_{50}$ is indicated at the bottom right. Error bars represent the standard deviation (SD) of the mean from independent replicates. FL = full-length; T = terminated product.

**(d)** Representative denaturing gel of time pausing assay. Transcription reactions were performed using 10 μM rNTPs. Scales were determined using increasing concentration of 2'-OMe-rNTPs. Positions of pause sites are indicated on the left. FL = full-length; T = terminated product; Ch = Chase reaction performed upon adding 500 μM rNTPs for 5 additional minutes at the end of the transcription reaction.

**(e)** Mapped reads from the *Bsu* data set published in Larson et al., Science 6187, 1042-1047 (2014).

**(f)** Representative denaturing gel showing termination and antitermination in the absence (-) and presence (+) of uncharged $tRNA^{Gly}$ in the context of the WT and MBox. Percentages of readthrough are indicated at the bottom of the gel. FL = full-length; T = terminated product.

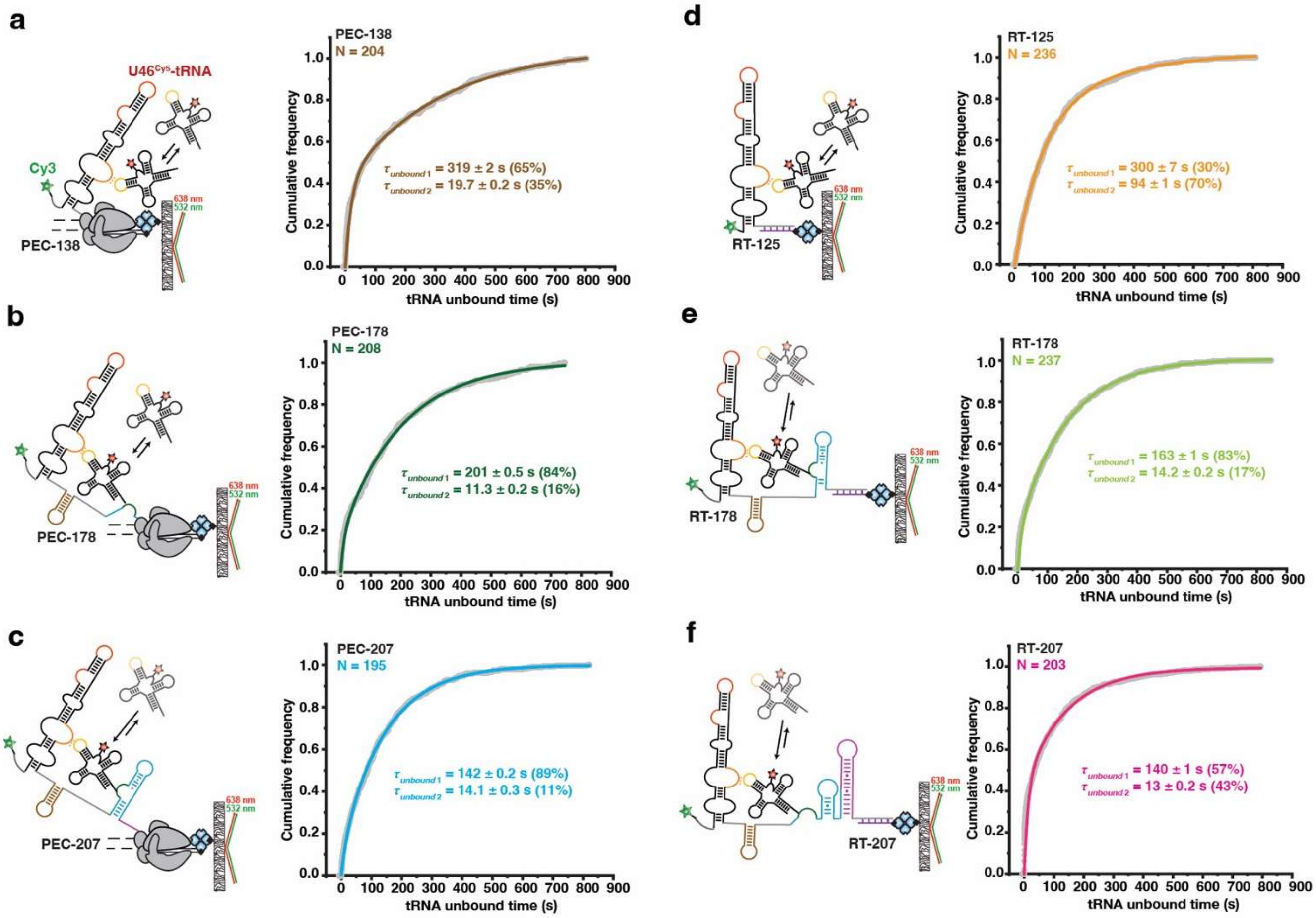


**Figure S2. Kinetics of U46$^{Cy5}$-tRNA recruitment to the *Bsu-GlyQS* T-Box riboswitch.**

Plots displaying the cumulative unbound dwell times of the U46$^{Cy5}$-tRNA to the *GlyQS*-Tbox riboswitch in the context of PEC-138 (a); PEC-178 (b); PEC-207 (c); RT-125 (d); RT-178 (e) and RT-207 (f). A cartoon depiction of the corresponding construct is indicated on the left of each plot. Total number of molecules analyzed for each construct is indicated.

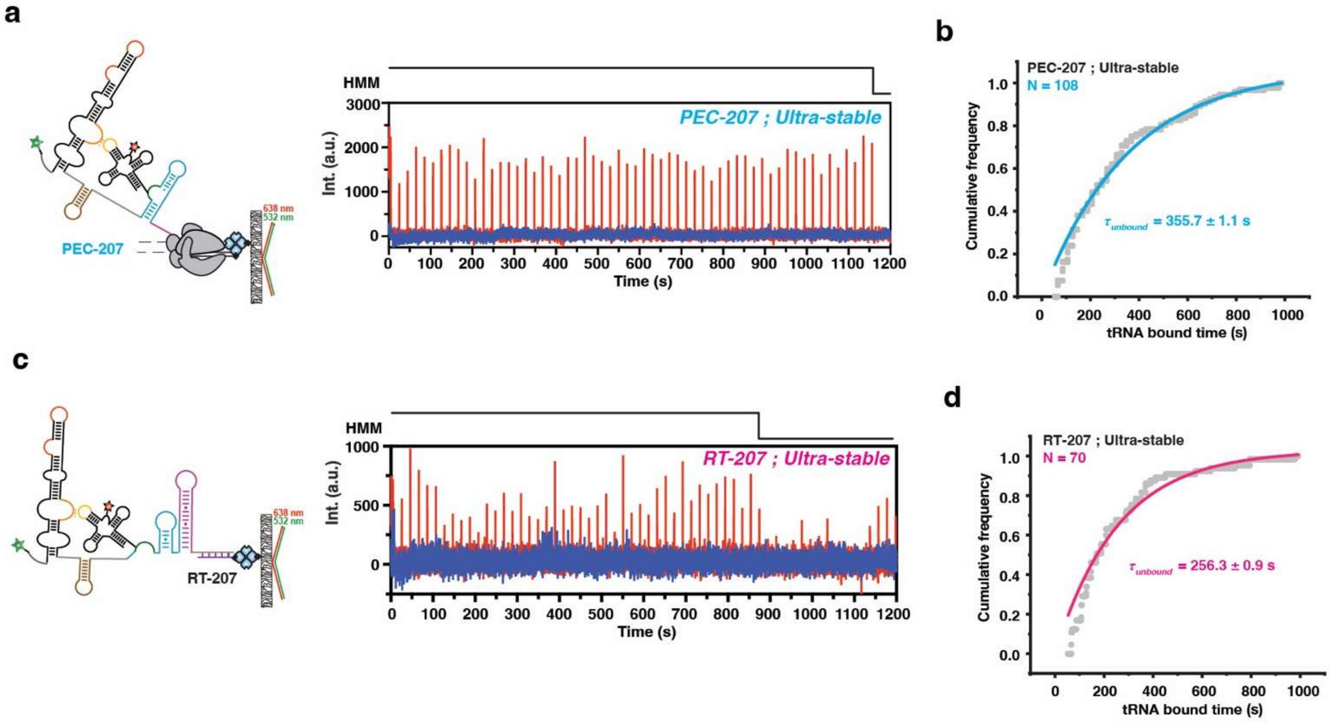


**Figure S3. The *Bsu-GlyQS* T-Box riboswitch can form ultra-stable complex with Gly-tRNA during transcription**

**(a)** Secondary structure of the *Bsu-GlyQS* T-box riboswitch in the context of the Paused Elongation Complexes (PEC) at the 207 position, alongside representative single-molecule trajectories showing U46$^{Cy5}$-tRNA ultra-stable binding (red) to the corresponding PEC (blue). Hidden Markov Modeling (HMM) is indicated on the top of the trace.

**(b)** Plot displaying the cumulative ultra-stable bound dwell times of U46$^{Cy5}$-tRNA in the PEC-207 construct. The overall bound ($\tau_{bound}$) time is indicated.

**(c)** Secondary structure of the *Bsu-GlyQS* T-box riboswitch in the context of the Released Transcript (RT) at the 207 position, alongside representative single-molecule trajectories showing U46$^{Cy5}$-tRNA ultra-stable binding (red) to the corresponding RT (blue). Hidden Markov Modeling (HMM) is indicated on the top of the trace.

**(d)** Plot displaying the cumulative ultra-stable bound dwell times of U46$^{Cy5}$-tRNA in the RT-207 construct. The overall bound ($\tau_{bound}$) time is indicated.

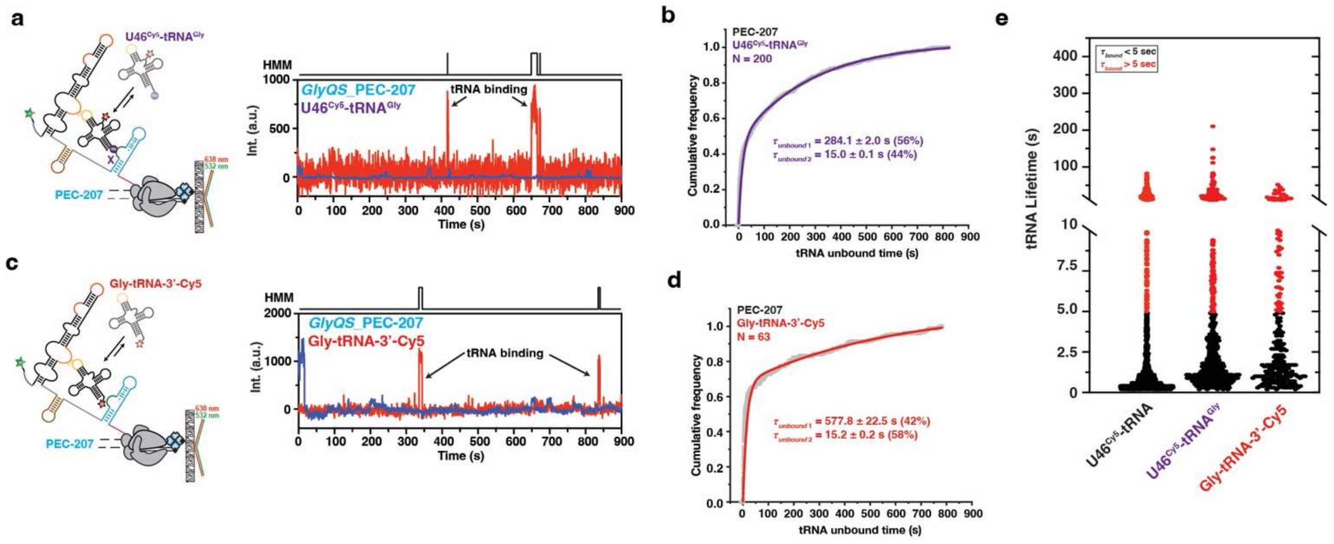


**Figure S4. Kinetic analysis of charged and 3'-Cy5 Gly-tRNA to the *Bsu-GlyQS* PEC-207**

**(a)** Secondary structure of the *Bsu-GlyQS* T-box riboswitch in the context of the Paused Elongation Complexes (PEC) at the 207 position, alongside representative single-molecule trajectories showing U46$^{Cy5}$-tRNA$^{Gly}$ binding (red) to the corresponding PEC (blue). Hidden Markov Modeling (HMM) is indicated on the top of the trace.

**(b)** Plot displaying the cumulative unbound dwell times of U46$^{Cy5}$-tRNA$^{Gly}$ in the PEC-207 construct. The overall unbound ($\tau_{unbound}$) times are indicated.

**(c)** Secondary structure of the *Bsu-GlyQS* T-box riboswitch in the context of the Paused Elongation Complexes (PEC) at the 207 position, alongside representative single-molecule trajectories showing Gly-tRNA-3'Cy5 binding (red) to the corresponding PEC (blue). Hidden Markov Modeling (HMM) is indicated on the top of the trace.

**(d)** Plot displaying the cumulative unbound dwell times of Gly-tRNA-3'Cy5 in the PEC-207 construct. The overall unbound ($\tau_{unbound}$) times are indicated.

**(e)** Plot displaying the overall distribution of tRNA lifetimes for the different fluorescently-labeled tRNAs tested. Short binding events ($\tau_{bound}$ <5 s) are indicated in black and long-lived binding events ($\tau_{bound}$ >5 s) are indicated in red. The total number of molecules analyzed is as follow: U46$^{Cy5}$-tRNA = 195; U46$^{Cy5}$-tRNA$^{Gly}$ = 200; Gly-tRNA-3'Cy5= 63.

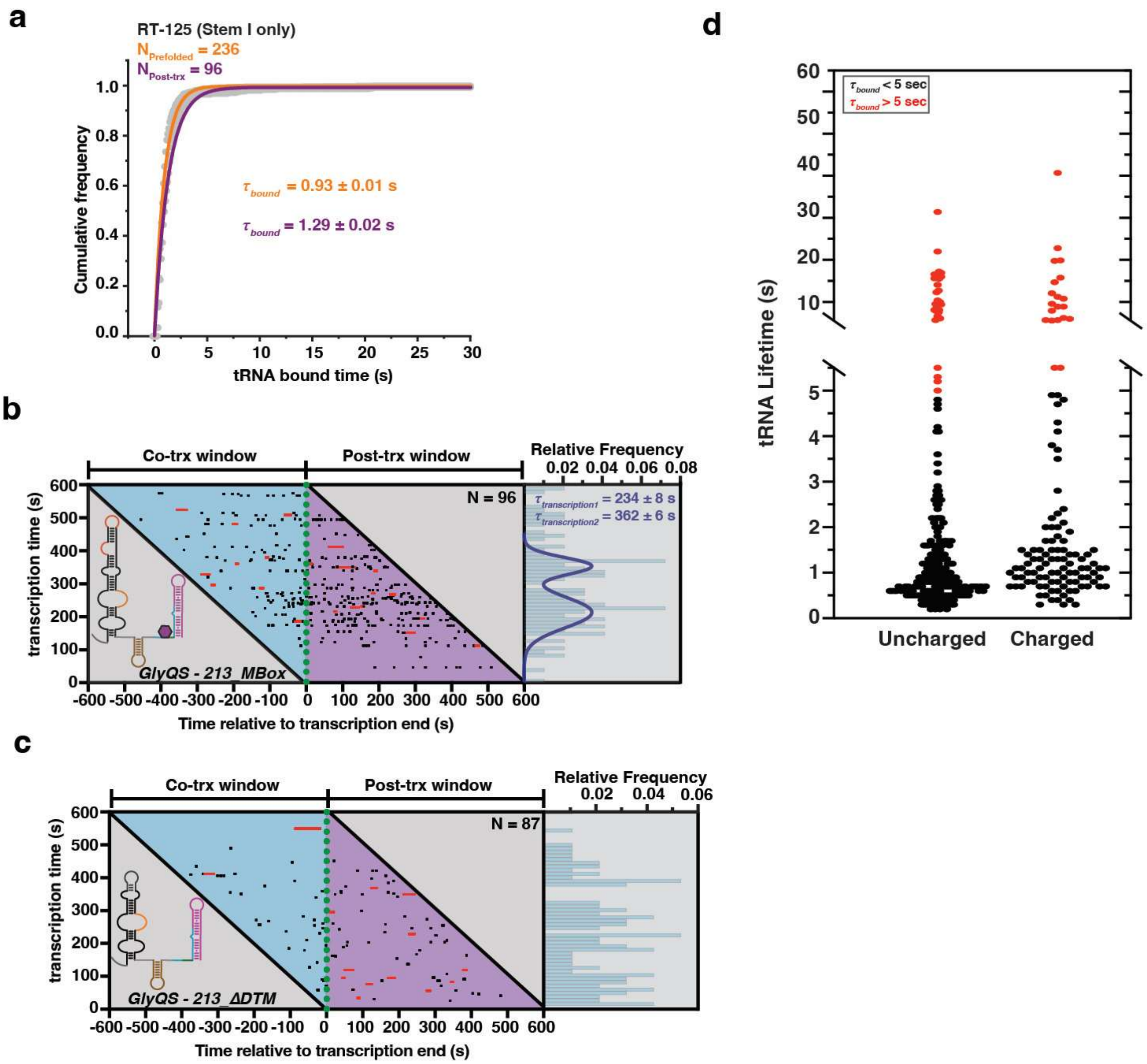


**Figure S5. Real-time analysis of tRNA binding during active transcription.**

**(a)** Plot displaying the cumulative bound dwell times of U46$^{Cy5}$-tRNA in the RT-125 construct pre-folded before single-molecule imaging (orange) or co-transcriptionally folded on the microscope slide (purple). The overall bound ($\tau_{bound}$) times are indicated. Total number of molecules analyzed for each construct is indicated.

**(b-c)** Rastergrams of U46$^{Cy5}$-tRNA binding (black and red bars) during (cyan background) and just after transcription (magenta background) of the *GlyQS*_213_Mbox (b) and *GlyQS*_213_ΔDTM (c) DNA templates; each horizontal line represents a single transcribed molecule. The lengths and colors of horizontal bars indicate the lifetimes of individual complexes. On the x axis, t = 0 represents the time relative to the transcription end marked by occurrence of the PIFE signal. Histogram of transcription time is indicated on the right. The indicated error is the error of the gaussian fit.

**(d)** Plot displaying the overall distribution of tRNA lifetimes for the uncharged and charged Gly-tRNA during transcription performed using 25 μM rNTPs. Short binding events ($\tau_{bound}$ <5 s) are indicated in black and long-lived binding events ($\tau_{bound}$ >5 s) are indicated in red. The total number of molecules analyzed is as follow: Uncharged = 181; Charged = 47.

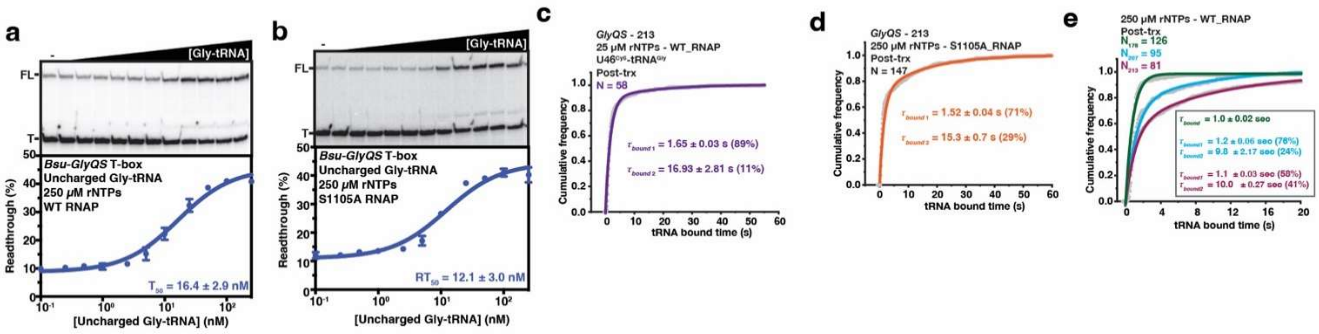


**Figure S6. Impact of the transcription on co-transcriptional tRNA sensing.**

**(a-b)** Representative denaturing gels (top) and plots of the fraction of transcription readthrough versus the concentration of Gly-tRNA (bottom) in the context of fast transcription (a) or slower transcription (b). Transcription reactions were performed using 250 µM rNTPs. The $RT_{50}$ is indicated at the bottom right. Error bars represent the standard deviation (SD) of the mean from independent replicates. FL = full-length; T = terminated product.

**(c)** Plot displaying the cumulative bound dwell times of U46$^{Cy5}$-tRNA$^{Gly}$ to *GlyQS-213* after real-time transcription completion using fast transcription regime. The overall bound ($\tau_{bound}$) times are indicated. Total number of molecules analyzed is indicated.

**(d)** Plots displaying the cumulative bound dwell times of U46$^{Cy5}$-tRNA to *GlyQS-213* after real-time transcription completion using slow transcription regime. The overall bound ($\tau_{bound}$) times are indicated. Total number of molecules analyzed is indicated.

**(e)** Plots displaying the cumulative bound dwell times of U46$^{Cy5}$-tRNA to *GlyQS-213; GlyQS-207* and *GlyQS-178* after real-time transcription completion using fast transcription regime. The overall bound ($\tau_{bound}$) times are indicated. Total number of molecules analyzed for each construct is indicated.

**Table S1: Oligonucleotides used in this study.**

| Oligonucleotide | Sequence (5'-3') |
|---|---|
| T7A1-PCR | TCCAGATCCCGAAAATTTATCAAAAAGAGTATTG |
| GlyQS-Tbox-FLrev | CAAATGTGGTATGGCTGATTCAAGGT |
| Tbox_GlyQS-RT125 | AGACCACGTTGAAAGATTGGGTTACCTCTTTTTAAAATTGCTCAAGAATGCCTTTCG |
| Tbox_GlyQS-EC178 | /5Biosg/AAACATAGGGACGAGAGTTCTCCC |
| Tbox_GlyQS-EC207 | /5Biosg/AAACTCCCGTCTCTATGCTTGCC |
| Tbox_GlyQS-RT178 | AGACCACGTTGAAAGATTGGGTTACGACGAGAGTTCTCCCGCGG |
| Anchor_bio | /5Biosg/AGACCACGTTGAAAGATTGGGTTAC |
| Tbox_GlyQS-RT207 | AGACCACGTTGAAAGATTGGGTTACTCTCTATGCTTGCCAGCCGC |
| GlyQS-Tbox-RT125_Cy3 | /5Cy3/CTCTTTTTAAAATTGCTCAAGAATGCCTTTCG |
| GlyQS-Tbox-RT213_Cy3 | /5Cy3/CTCCCGTCTCTATGCTTGCCA |
| GlyQS-Tbox-EC138 | /5BiosG/TGCTGAGAACAAAATCCCAGCCT |

| | |
|---|---|
| GlyQS-Tbox-mutTbox (2) | TTCTCCCGCGGTAGGTCCCTAGTTGCTG |
| GlyQS-Tbox-mutTbox (3) | CAGCAACTAGGGACCTACCGCGGGAGAA |
| GlyQS-DeltaDTM (2) | GAATGCCTTTCGTTGCGTGCCGCCGAAGCGGGTAAACGCAAGTACTTCTTTC |
| GlyQS-DeltaDTM (3) | GAAAGAAGTACTTGCGTTTACCCGCTTCGGCGGCACGCAACGAAAGGCATTC |
| GlyQS-178_Cy3 | /5Cy3/GACGAGAGTTCTCCCGCGGT |
| GlyQS-RT207_Cy3 | /5Cy3/TCTCTATGCTTGCCAGCCGC |
| Tbox-IntAmino-3Cy3 | /5Cy3/CTTTCATAAGTTGACTGCGGCAGCAACCAAAAAACTCCCGT/iAmMC6T/CTCTATGCTTGCCAGCCGC |
| Tbox-MutAT (2) | CCAGCCGCAAACATACCCACGAGAGTTCTC |
| Tbox-MutAT (3) | GAGAACTCTCGTGGGTATGTTTGCGGCTGG |
| GlyQS-MutAT-comp(2) | CCCGCGGTTCCAGGGATGTTGCTGAGAAC |
| GlyQS-MutAT-comp(3) | GTTCTCAGCAACATCCCTGGAACCGCGGG |

**Table S2: tRNA binding kinetics on immobilized transcripts and Paused Elongation Complexes.**

| Construct | $k_{on}$ ($10^6$ $M^{-1}$ $s^{-1}$) | $k_{off}$ ($s^{-1}$) |
|---|---|---|
| **PEC-138** | Fast[a]: 8.12 ± 0.08 (35%)<br>Slow[a]: 0.50 ± 0.003 (65%)<br>Overall[b]: 0.75 | Fast[a]: 0.83 ± 0.01 (90%)<br>Slow[a]: 0.07 ± 0.01 (10%)<br>Overall[b]: 0.40 |
| **PEC-178** | Fast[a]: 14.2 ± 0.25 (16%)<br>Slow[a]: 0.80 ± 0.001 (84%)<br>Overall[b]: 0.94 | Fast[a]: 0.71 ± 0.02 (88%)<br>Slow[a]: 0.05 ± 0.006 (12%)<br>Overall[b]: 0.28 |
| **PEC-207_WT** | Fast[a]: 11.3 ± 0.2 (11%)<br>Slow[a]: 1.12 ± 0.001 (89%)<br>Overall[b]: 1.25 | Fast[a]: 0.81 ± 0.02 (75%)<br>Slow[a]: 0.05 ± 0.003 (25%)<br>Overall[b]: 0.17 |
| **PEC-207_Mbox** | Fast[a]: NA<br>Slow[a]: 0.70 ± 0.001<br>Overall[b]: 0.70 | Fast[a]: 1.16 ± 0.01<br>Slow[a]: NA<br>Overall[b]: 1.16 |
| **PEC-207_MAT** | Fast[a]: NA<br>Slow[a]: 0.69 ± 0.008<br>Overall[b]: 0.69 | Fast[a]: 1.13 ± 0.01<br>Slow[a]: NA<br>Overall[b]: 1.13 |
| **PEC-207_MATcp** | Fast[a]: 7.49 ± 0.6 (40%)<br>Slow[a]: 0.59 ± 0.003 (60%)<br>Overall[b]: 0.92 | Fast[a]: 0.76 ± 0.005 (61%)<br>Slow[a]: 0.09± 0.001 (37%)<br>Overall[b]: 0.20 |
| **RT-125** | Fast[a]: 1.70 ± 0.02 (70%)<br>Slow[a]: 0.53 ± 0.01 (30%)<br>Overall[b]: 1.03 | Fast[a]: 1.08 ± 0.01<br>Slow[a]: NA<br>Overall[b]: 1.08 |
| **RT-125_ΔDTM** | Fast[a]: NA<br>Slow[a]: 0.70 ± 0.003<br>Overall[b]: 0.70 | Fast[a]: 1.08 ± 0.02 (89%)<br>Slow[a]: 0.12 ± 0.01 (10%)<br>Overall[b]: 0.60 |
| **RT-178** | Fast[a]: 11.27± 0.16 (17%)<br>Slow[a]: 0.98 ± 0.01 (83%)<br>Overall[b]: 1.16 | Fast[a]: NA<br>Slow[a]: 0.27 ± 0.004<br>Overall[b]: 0.27 |
| **RT-207** | Fast[a]: 12.3 ± 0.19 (43%)<br>Slow[a]: 1.14 ± 0.008 (57%)<br>Overall[b]: 1.55 | Fast[a]: 0.53 ± 0.009 (75%)<br>Slow[a]: NA<br>Overall[b]: 0.53 |
| **PEC-207_WT + Charged tRNA** | Fast[a]: 5.33 ± 0.04 (44%)<br>Slow[a]: 0.28 ± 0.02 (56%)<br>Overall[b]: 0.48 | Fast[a]: 0.26 ± 0.004 (92%)<br>Slow[a]: 0.02 ± 0.003 (8%)<br>Overall[b]: 0.14 |
| **PEC-207_WT + 3'-Cy5-tRNA** | Fast[a]: 5.26 ± 0.07 (58%)<br>Slow[a]: 0.14 ± 0.005 (42%)<br>Overall[b]: 0.32 | Fast[a]: 0.47 ± 0.01 (68%)<br>Slow[a]: 0.09 ± 0.004 (31%)<br>Overall[b]: 0.21 |

[a]Values were calculated from single or double-exponential fits of the pool data from all the experiments in a given condition. The percentages indicate the contribution of each phase to the overall rate constant. The reported error is the standard deviation of the fit. In the case of single-exponential fit, only one value is reported arbitrarily relative to the rate constant.

[b]Values represent the weight average taking into consideration the proportions of each dwell time.

**Table S3: tRNA bound times kinetics observed during real-time transcription.**

| Construct | rNTPs | RNAP | Co-Trx ($s^{-1}$) | Post-Trx ($s^{-1}$) |
|---|---|---|---|---|
| **RT-125** | 25 µM | WT | Fast[a]: 0.77 ± 0.02<br>Slow[a]: NA<br>Overall[b]: 0.77 | Fast[a]: 0.78 ± 0.01<br>Slow[a]: NA<br>Overall[b]: 0.78 |
| **RT-213_WT** | 25 µM | WT | Fast[a]: 0.92 ± 0.03 (84%)<br>Slow[a]: 0.07 ± 0.02 (16%)<br>Overall[b]: 0.32 | Fast[a]: 0.69 ± 0.01 (61%)<br>Slow[a]: 0.08 ± 0.02 (39%)<br>Overall[b]: 0.17 |
| **RT-213_WT – Charged tRNA** | 25 µM | WT | Fast[a]: NA<br>Slow[a]: 0.27 ± 0.004<br>Overall[b]: 0.27 | Fast[a]: 0.60 ± 0.01 (89%)<br>Slow[a]: 0.06 ± 0.01 (11%)<br>Overall[b]: 0.30 |
| **RT-213_WT** | 250 µM | WT | Fast[a]: 0.98 ± 0.04 (78%)<br>Slow[a]: 0.03 ± 0.01 (22%)<br>Overall[b]: 0.13 | Fast[a]: 0.90 ± 0.02 (58%)<br>Slow[a]: 0.1 ± 0.002 (41%)<br>Overall[b]: 0.21 |
| **RT-213_WT** | 250 µM | S1105A | Fast[a]: 0.99 ± 0.03 (84%)<br>Slow[a]: 0.03 ± 0.01 (16%)<br>Overall[b]: 0.18 | Fast[a]: 0.66 ± 0.02 (71%)<br>Slow[a]: 0.07 ± 0.003 (29%)<br>Overall[b]: 0.18 |
| **RT-213_Mbox** | 25 µM | WT | Fast[a]: 0.89 ± 0.02<br>Slow[a]: NA<br>Overall[b]: 0.89 | Fast[a]: 0.88 ± 0.02<br>Slow[a]: NA<br>Overall[b]: 0.88 |
| **RT-213_ΔDTM** | 25 µM | WT | Fast[a]: 0.58 ± 0.01<br>Slow[a]: NA<br>Overall[b]: 0.58 | Fast[a]: 0.78 ± 0.02 (82%)<br>Slow[a]: 0.05 ± 0.01 (18%)<br>Overall[b]: 0.21 |
| **RT-178_WT** | 250 µM | WT | Fast[a]: 0.76 ± 0.06<br>Slow[a]: NA<br>Overall[b]: 0.76 | Fast[a]: 1.0 ± 0.02<br>Slow[a]: NA<br>Overall[b]: 1.0 |
| **RT-207_WT** | 250 µM | WT | Fast[a]: 0.60 ± 0.03<br>Slow[a]: NA<br>Overall[b]: 0.60 | Fast[a]: 0.83 ± 0.04 (76%)<br>Slow[a]: 0.1 ± 0.02 (24%)<br>Overall[b]: 0.30 |

[a]Values were calculated from single or double-exponential fits of the pool data from all the experiments in a given condition. The percentages indicate the contribution of each phase to the overall rate constant. The reported error is the standard deviation of the fit. In the case of single-exponential fit, only one value is reported arbitrarily relative to the rate constant.

[b]Values represent the weight average taking into consideration the proportions of each dwell time.
